# ‘Low’ LRs obtained from DNA mixtures: On calibration and discrimination performance of probabilistic genotyping software

**DOI:** 10.1101/2024.06.06.597689

**Authors:** M. McCarthy-Allen, Ø. Bleka, R. Ypma, P. Gill, C. Benschop

**Affiliations:** Netherlands Forensic Institute, Division of Biological Traces; Oslo University Hospital, Department of Forensic Sciences; Netherlands Forensic Institute, Division of Digital and Biometric Traces; University of Oslo, Institute of Clinical Medicine, Department of Forensic Medicine

**Keywords:** Forensic DNA, DNA mixtures, probabilistic genotyping, calibration, DNAStatistX, EuroForMix

## Abstract

The validity of a probabilistic genotyping (PG) system is typically demonstrated by following international guidelines for the developmental and internal validation of PG software. These guidelines mainly focus on discriminatory power. Very few studies have reported with metrics that depend on calibration of likelihood ratio (LR) systems. In this study, discriminatory power as well as various calibration metrics, such as Empirical Cross-Entropy (ECE) plots, pool adjacent violator (PAV) plots, log likelihood ratio cost (Cllr and Cllr*^cal^*), fiducial calibration discrepancy plots, and Turing’ expectation were examined using the publicly-available PROVEDIt dataset. The aim was to gain deeper insight into the performance of a variety of PG software in the ‘lower’ LR ranges (∼LR 1-10,000), with focus on DNAStatistX and EuroForMix which use maximum likelihood estimation (MLE). This may be a driving force for the end users to reconsider current LR thresholds for reporting. In previous studies, overstated ‘low’ LRs were observed for these PG software. However, applying (arbitrarily) high LR thresholds for reporting wastes relevant evidential value. This study demonstrates, based on calibration performance, that previously reported LR thresholds can be lowered or even discarded. Considering LRs >1, there was no evidence for miscalibration performance above LR ∼1,000 when using Fst 0.01. Below this LR value, miscalibration was observed. Calibration performance generally improved with the use of Fst 0.03, but the extent of this was dependent on the dataset: results ranged from miscalibration up to LR ∼100 to no evidence of miscalibration alike PG software using different methods to model peak height, HMC and STRmix.

This study demonstrates that practitioners using MLE-based models should be careful when low LR ranges are reported, though applying arbitrarily high LR thresholds is discouraged. This study also highlights various calibration metrics that are useful in understanding the performance of a PG system.

**Highlights:**

- Discriminatory power and calibration performance of PG software are evaluated.
- The utility of various calibration metrics are explored in ‘low’ LR ranges.
- Focus was on DNAStatistX and EuroForMix software using the MLE method.
- Calibration performance was dependent on Fst value and dataset size.
- Results suggest reconsideration of lower LR thresholds and cautious reporting of ‘low’ LRs.

## 1. Introduction

### 1.1 Evaluating performance of PG software

Forensic DNA mixture interpretation is aided by probabilistic genotyping (PG) software and is currently used in many, if not most, forensic DNA laboratories worldwide. These PG software provide weights of evidence (likelihood ratios; LR values) for potential contributors in single source up to very complex mixed DNA profiles. Application of PG software in forensic DNA casework is preceded by its validation. Various guidelines on validation of PG systems have been reported [1, 2, 3, 4, 5, 6] and split into developmental and internal validation.

Developmental validation is undertaken by the developers and includes aspects such as ‘demonstrating that the model is an acceptable approach’ (e.g. through peer review publication), ‘demonstrating the performance of the models in cases where the true state is known’ (i.e. sensitivity, specificity, precision), and ‘demonstrating that the calculations made by the software emulating the model are correct when the ‘true’ state is not known’ [5]. Besides the technical aspects, developmental validation includes aspects such as availability of a user manual – including version numbers and changes – and access to training resources.

Internal validation is conducted by the forensic science end users of the software to ascertain that it is fit for purpose, under the conditions that it is intended to be used [5]. This also includes developing standard operating procedures (SOPs), outlining the types of cases and data that can be analysed.

SOPs for DNA mixture interpretation vary between laboratories. There are notable differences in the criteria to determine what is too complex to interpret [7] and which LR values to report. With respect to complexity, some laboratories may limit LR calculations to mixtures with ≤3 (estimated) contributors, whilst other laboratories may extend this (e.g. to 5), or use a ‘top-down’ approach for mixtures with >4 estimated contributors. Furthermore, some laboratories limit LR calculations based on the maximum number of unseen alleles for the person of interest. In terms of LR values reported, many laboratories apply a minimum reporting threshold, which may be an LR of e.g. 10, 100, 1,000, 10,000, [8] or even a million [9].

However, the use of a reporting threshold has been argued against (see e.g. [10, 11]). Furthermore, it must be noted that LR thresholds are not the only factor that affect whether a laboratory reports a weight of evidence. For example, an LR minimum threshold of e.g. 10,000 versus 1,000 seems large. However, if a laboratory that applies an LR threshold of 1,000 uses a qualitative LR model, an STR typing system with 15 autosomal markers, and/or an Fst correction of 0.03, whilst the laboratory with an LR threshold of 10,000 uses a quantitative model, an STR typing system with 23 autosomal markers and/or an Fst correction of 0.01, the actual differences in LR may not be that different. This point is further exemplified by comparing SOP decisions of hypothetical laboratories, A and B. Laboratory A has no lower LR threshold for reporting, but does not calculate an LR due to profile complexity. Laboratory B does compute an LR with the same complex mixture profile, however, they have a lower LR reporting threshold. In turn, both A and B may report no LR, despite different SOPs. Therefore, interlaboratory comparisons of LR thresholds are not very informative because they do not solely dictate whether a laboratory reports on a sample. Nevertheless, examining the performance of PG software, specifically in the ‘lower’ LR ranges, and under various conditions is a worthy consideration, since it may be a driving force to persuade end users to revisit current reporting thresholds. If a PG software is perfectly calibrated throughout the entire LR range, a lower LR threshold should not be needed.

Prior to examining LRs in the lower ranges, it is important to note that low LRs (e.g. LR=100) are not ‘less reliable’ than high LRs (e.g. LR=10^6^) provided that the PG model is well calibrated. I.e. it can be demonstrated that the behaviour of LRs follows mathematically derived expectations. Furthermore, as described in [12], a complexity threshold should not be taken as evidence of the non-reliability of a PG model’s performance, but rather a practical or business decision.

In this study, we examine and discuss *discriminatory power* and *calibration* performance of PG software. We focus on the ‘lower range’ LRs of DNAStatistX [8]. In particular, on LRs below 10,000, constituting a minimum threshold for reporting LRs as presented in [8]. Furthermore, we compare to EuroForMix [13] LRs in this range. Both PG software employ the same maximum likelihood estimate (MLE) method for parameter estimation, and have been observed to overstate ‘lower range’ LRs relative to other PG software. Specifically, previous studies using DNAStatistX /EuroForMix have shown that:

(i) the LRs for H_d_-true scenarios tend towards neutral evidence if the number of contributors (NoC) is over-assigned [14]; and;
(ii) the LRs for H_d_-true scenarios, although <1, can be much larger (i.e. tend towards neutral evidence) than LRs obtained using other PG software (e.g. [15, 16, 17, 18]).

On the other hand, H_p_-true LRs in the ‘higher ranges’, have presented strong similarities to other PG software, as long as the same input data and propositions were used (e.g. [15, 19, 20, 21]).

Discriminatory power [22] in this context refers to a model’s ability to distinguish between persons of interest (POIs) being true-contributors or non-contributors to DNA profiles. Here, false negatives, false positives, sensitivity, specificity, accuracy, positive predictive value (PPV), and negative predictive value (NPV) can be assessed. These can be presented in, for instance, receiver operating characteristic (ROC) plots and the associated area under the ROC plot (AUC) [15, 17, 20, 23], scatter or LR distribution plots [8, 14, 15, 17, 23], or Accuracy/Misleading Evidence tables [15, 17, 20, 23]. These metrics for DNAStatistX and/or EuroForMix have been reported in previous studies.

Calibration refers to whether the LRs assigned by a model follow the mathematical properties that these LRs should have, i.e. whether the proportion of times we observe an LR of *x* under H_p_ is *x* times higher than under H_d_. For example, a model that always assigns an LR of 1 is perfectly calibrated, but has no discriminatory power. Different metrics have been proposed to assess calibration, which vary in use across different forensic disciplines [24, 25, 26]. Examples are Tippett plots [27], Turing tests or non-contributor performance tests [28], calibration tables [18, 29], Empirical Cross-Entropy (ECE) plots [27, 30, 31], log likelihood ratio cost (Cllr and Cllr*^cal^*) [30, 32], pool adjacent violators (PAV) plots, devPav [30] and fiducial calibration discrepancy plots [31]. However, except for Tippett plots and calibration tables, very few studies have reported on these metrics for PG systems [26].

In the current study, we employ several of these metrics to assess the discriminatory power and calibration performance of PG software. These metrics are explained in the following section. As mentioned, we focus on DNAStatistX and EuroForMix results in LR ranges <10,000. The results obtained in this study are further compared to the STRmix and HMC results obtained in the study of Susik & Sbalzarini [17]. Ultimately, we aim to interrogate, and make recommendations with respect to reporting lower range LRs obtained using the MLE method that is incorporated into DNAStatistX and EuroForMix.

### 1.2 Calibration performance metrics

Various metrics exist to assess calibration performance of LR based systems. In this section, calibration tables, fiducial calibration discrepancy plots, PAV plots (with devPAV values), Empirical Cross-Entropy plots (with Cllr and Cllr*^cal^*values) and Turing tests are further elucidated.

#### 1.2.1 Calibration tables

Calibration tables can be created as described in e.g. Bright et al. [29]. To create such tables, raw counts are needed of true (H_p_-true) and false (H_d_-true) contributor scenarios whose posterior probabilities (based on Bayes Theorem) fall into the defined ranges. From this, observed relative frequencies of H_p_-true scenarios can be calculated within each posterior probability range as 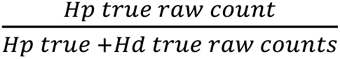. Good calibration is demonstrated by the observed relative frequency values falling within their respective posterior probability range.

#### 1.2.2 Fiducial calibration discrepancy plots

Fiducial calibration discrepancy plots provide information for regions of LR values (intervals) on whether evidence is overstated and/ or understated. Furthermore, they provide a magnitude and direction of discrepancy between reported LRs and the empirically supported values for well-calibrated LRs, and provide an approach for obtaining confidence bands for the calibration discrepancy [31].

As in the example (Fig. 1), log_10_LR intervals are plotted against interval-specific calibration discrepancy. The red line represents perfect calibration (i.e. no discrepancy from empirically supported values for well-calibrated LRs). The dark blue line represents the point estimate of calibration discrepancy for a given interval. This point estimate is calculated as the median of the fiducial distribution. The black lines (95% pointwise confidence intervals) and cyan lines (95% simultaneous confidence bounds) indicate uncertainty in the calibration discrepancy estimates. The black lines are used when assessing the calibration discrepancy of a single LR, whereas the cyan lines are used when assessing the calibration discrepancy of the overall LR system.

**Figure 1.**
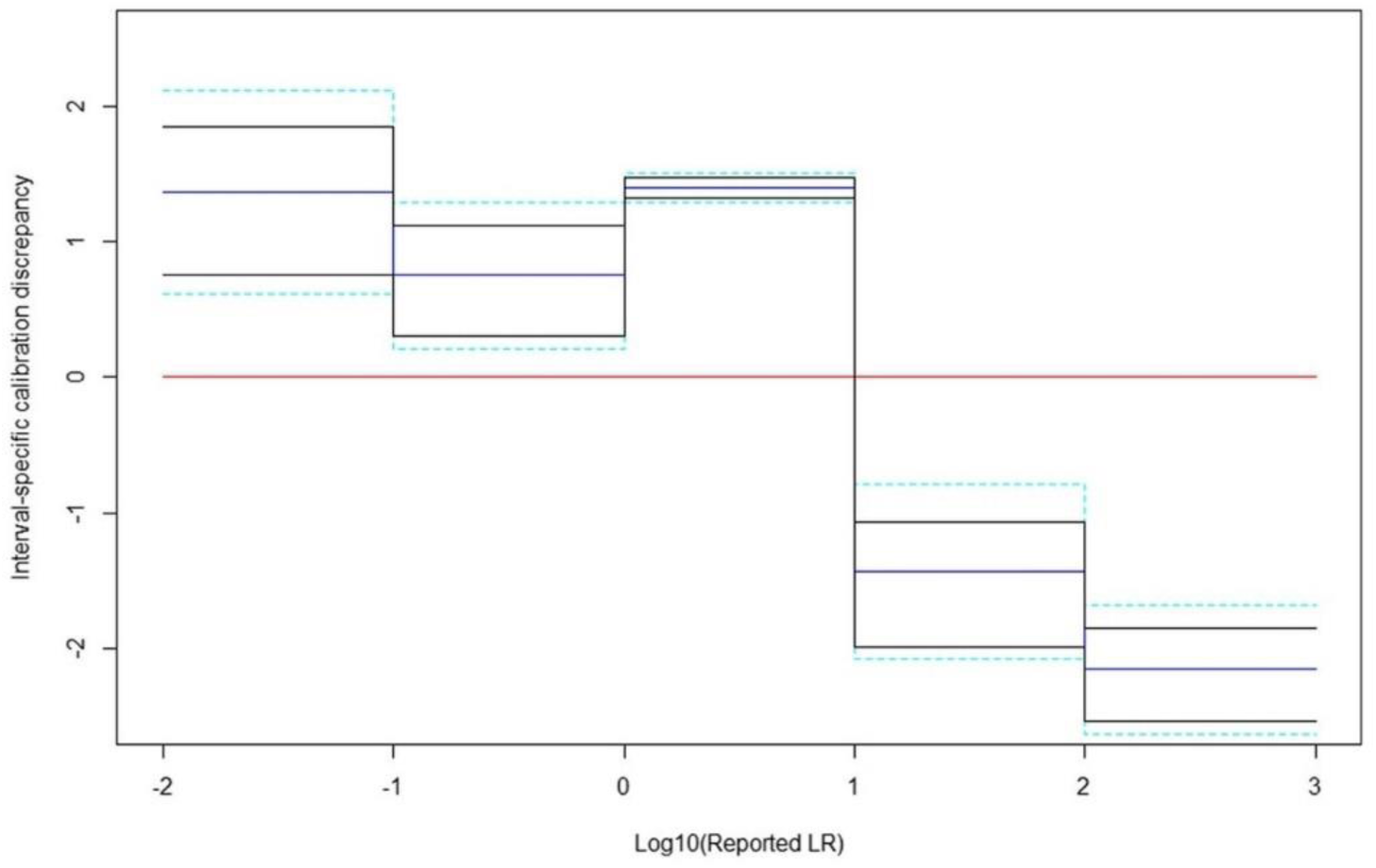
Fiducial calibration discrepancy plot example from [31].

Good calibration is indicated by tight confidence bounds that encompass the red line. Poor calibration is indicated by the red line sitting outside the confidence bounds. If the red line is above or below the bounds, the LRs are understated or overstated respectively, by a factor that may be read from the plot. For instance, Fig. 1 shows that LRs between 10^2^-10^3^ are overstating the evidence by a factor of at least 10^2^.

#### 1.2.3 Pool adjacent violator (PAV) plots with devPAV values

Like fiducial calibration discrepancy plots, PAV plots also indicate the magnitude and direction of calibration discrepancy, but taking empirically-defined, rather than user-defined bins of LRs. Generally, if we want to improve the calibration properties of a system without altering its discriminative properties, we can apply a non-decreasing transformation to it. The PAV algorithm applies a non-decreasing transformation to the LR values, such that it minimizes the sum of squared errors (i.e. isotonic regression). In step-wise adjustments, the algorithm pools adjacent LRs that violate the monotonicity constraint and then replaces these values with their mean. This is repeated throughout the sequence of LRs until no monotonicity constraint violators remain [32]. The algorithm is typically not used during the development of an LR system as it tends to overfit the data. This makes it a stringent test of calibration properties – if the PAV algorithm does not alter the LR values on a dataset, we can conclude the system was well calibrated (on these data).

In a PAV plot, the PAV-transformed log_10_LRs (post-calibrated) are plotted against the original log_10_LRs (pre-calibrated) (i.e. a ‘PAV transform’), with an x=y line (identity line) (Fig. 2). The deviation between the PAV transform and identity lines indicates how well the original set of LRs were calibrated. For a well-calibrated set of LRs, the PAV transform follows the identity line. Poor calibration is indicated by large deviations. If the LRs are originally understated, the PAV transform line lies to the left of the identity line. If the LRs are originally overstated, the PAV transform line lies to the right of the identity line [30].

**Figure 2.**
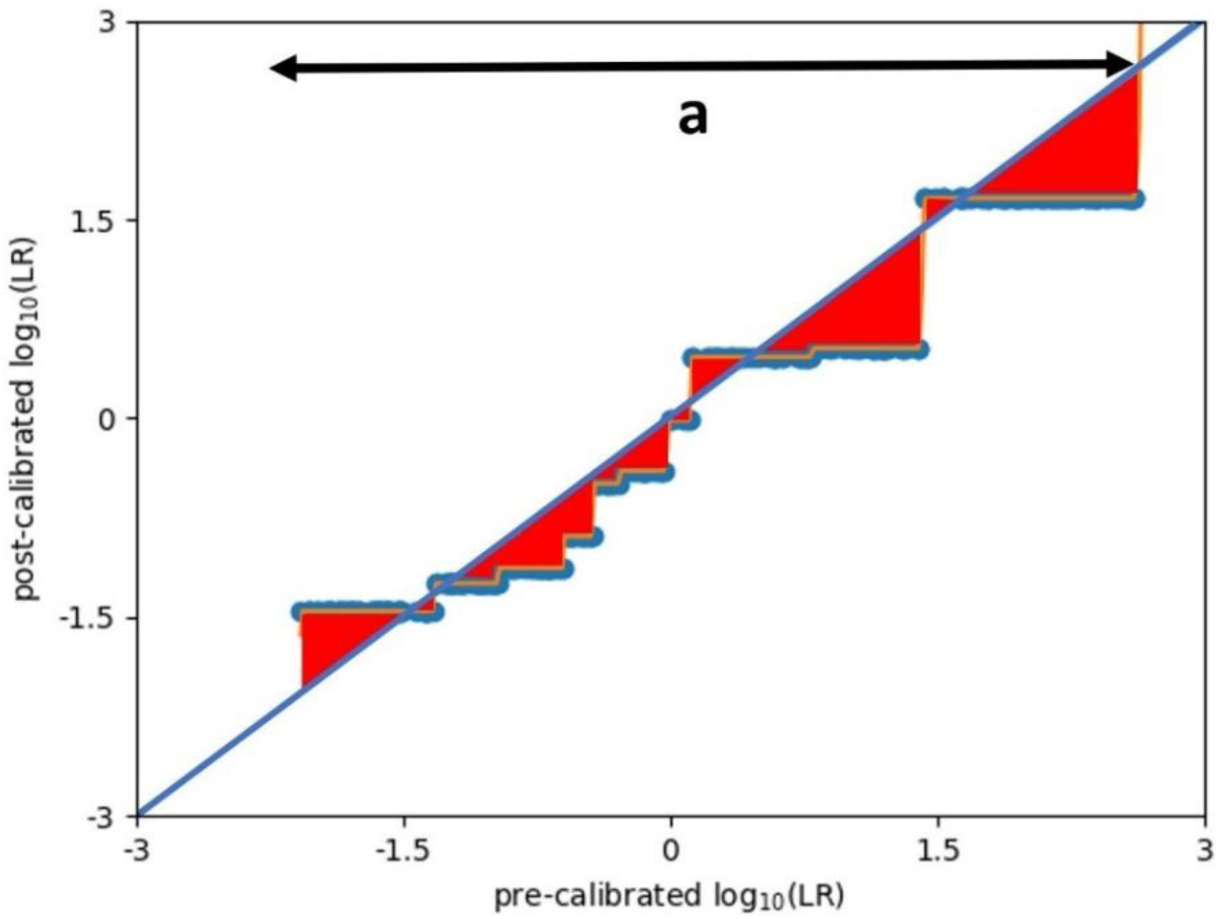
PAV plot example with devPAV calculation, where devPAV is equal to the area shaded in red divided by the range of the PAV transform indicated by *a*. Calibration here is okay as the diagonal is followed, but not perfect as seen from the occasional deviation from the diagonal.

Vergeer et al. propose ‘devPAV’ as a calibration metric, defined as the “average absolute deviation of the PAV-transform to the identity line” [30]. The metric is calculated as in Fig. 2, by dividing the area between the PAV transform and identity lines (red shaded) by the finite range of the PAV transform (line marked a). Good calibration is indicated by a small devPAV value.

#### 1.2.4 Empirical Cross-Entropy (ECE) plots with Cllr and Cllr*^cal^* values

LR performance can also be assessed using Empirical Cross-Entropy (ECE) plots. ECE is a concept from information theory, and can be thought of as the mean information needed to know the ground truth after considering the LR. It is a measure of accuracy, as a combination of discriminatory power and calibration [27]. ECE plots show model performance as a function of the prior probability. In this respect they differ from calibration tables, fiducial calibration discrepancy plots and PAV plots, which focus on the calibration as a function of the LR value. ECE plots, however, do not provide information on the extent to which evidence is being understated or overstated like fiducial calibration discrepancy plots and PAV plots.

ECE plots are made of three curves (Fig. 3):

(*i*) *ECE curve (solid orange):* represents ECE of all the LRs in a dataset, as a function of the prior odds. Smaller ECE values indicate better performance of the LR model.
(*ii*) *ECE curve after PAV (dashed green):* represents the ECE as a function of the prior odds, after transformation by the PAV-algorithm (as described in 1.2.3). The PAV-algorithm transforms the posterior probabilities in a dataset (calculated from the LRs) into an optimally calibrated set. The effect of the calibration component of ECE is minimized, leaving the effect of discriminatory power.
(*iii*) *“Neutral method” reference curve (dashed blue):* represents the ECE for a system that always outputs LR = 1 (i.e. neutral evidence) as a function of the prior odds (so when posterior odds = prior odds). At regions of the prior odds where the *ECE curve* is higher than the *“Neutral method” reference curve*, the LR model performs worse than if there was no evidence in support of either hypothesis. Such a model would make decisions worse than using no evidence at all. It indicates the presence of misleading LRs with respect to the ground truth state.

**Figure 3.**
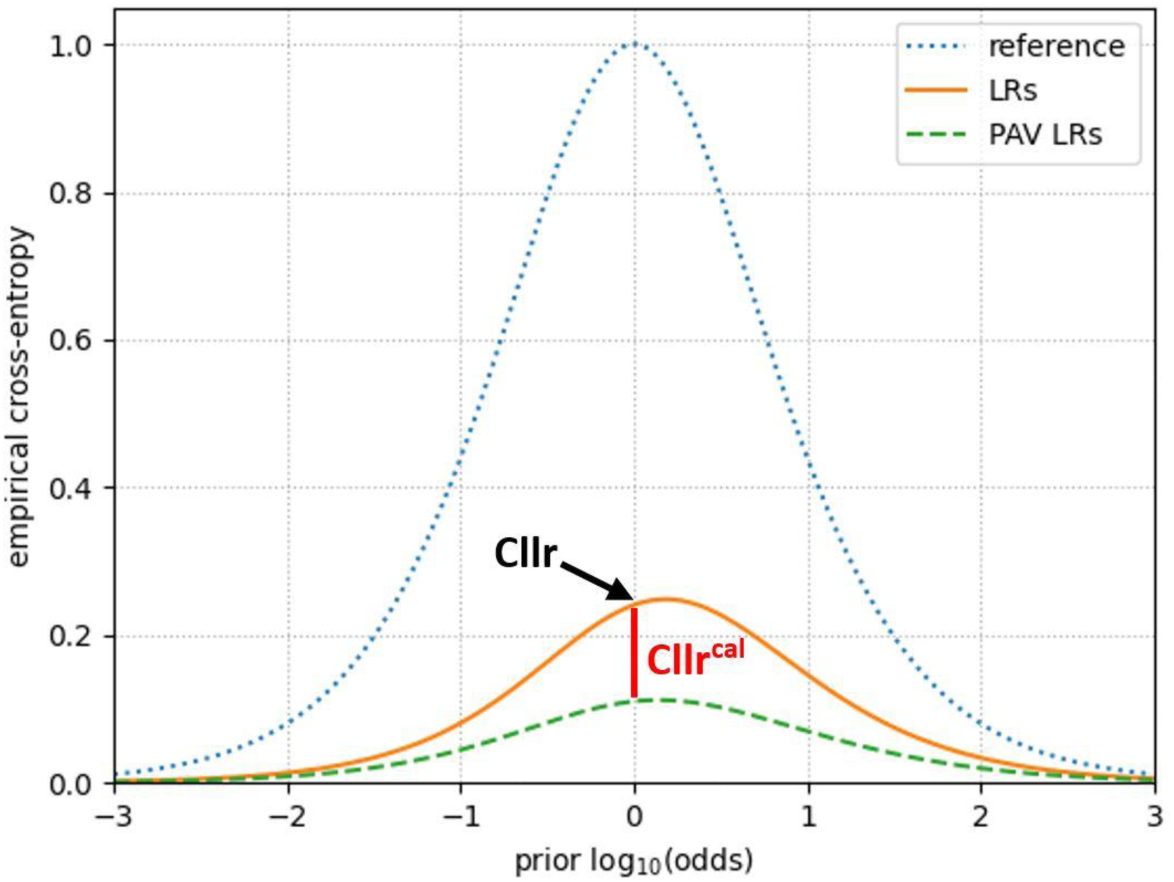
Example ECE plot with visual description of Cllr (black) and Cllr*^cal^* (red) calculation.

From ECE plots, one can assess the performance of an LR model with respect to:

- The performance of the LR model method compared to the *“Neutral method”.* The LR model performance is higher for values of the prior odds when the solid orange curve is lower than the dashed blue curve.
- *Discriminatory power*: a lower dashed green curve indicates the LRs are more discriminatory.
- *Calibration*: the closer the solid orange and dashed green curves, the better the calibration [27].

The ECE plot is often summarized by taking the *ECE curve* value at equal prior odds, called the log likelihood ratio cost (Cllr). In many forensic fields this is used as the single number most suited to measure LR system performance, as it takes into account both discriminatory power and calibration. The smaller the Cllr value, the better the performance [33]. Cllr*^cal^* is defined as the difference between Cllr and the Cllr obtained after PAV transformation, equivalent to the straight line distance between the *ECE curve* and *ECE curve after PAV* at a prior log_10_ odds of 0 (as in Fig. 3) [30]. Good calibration is indicated by a small Cllr*^cal^*value.

#### 1.2.5 Turing expectation tests (i.e. non-contributor tests)

Turing expectation tests, or non-contributor tests, are exploratory tests and can be used to investigate the robustness of the LR [34]. The general principle behind this is the same as in Tippet plots [30], that a good probabilistic model should be able to differentiate between true– and non-contributor scenarios. Therefore, this test also relates to the model’s *discriminatory power*.

With these tests, the LR*_x_* of a true-contributor scenario is compared to N LRs computed under the same model parameters, but with random non-contributors set as the POI. N is usually large and thus a distribution of random, non-contributor LRs is established.

The Turing expectation is used to compare non-contributor LRs to LR_*x*_, and in turn, evaluate model calibration and the robustness of LR_*x*_. This expectation can be summarised as: “the fraction of non-contributor scenarios producing an LR ≥ LR_*x*_ is expected to be *at most* 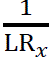. Therefore in N non-contributor tests, we expect LRs to be ≥ LR_*x*_ at most 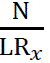 times” [12, 34, 35]. For example, if LR_*x*_ = 1,000, and we carry out N = 1,000 non-contributor tests, the non-contributor LRs should be ≥ LR_*x*_ at most 1 time. If this is true, the Turing expectation has passed. Failing Turing expectation indicates that a model is not well-calibrated. It has been suggested that these tests can be used in an exploratory manner on a case-by case basis to decide whether to report a (low) LR [34].

#### 1.2.6 Notes on terminology and data to asses calibration performance

At this point, a note on terminology is important. In the wider forensic science literature, ‘calibration’ of an LR system can refer to two distinct concepts: the degree to which the LRs it outputs adhere to the mathematical property of calibration, and the process used to optimise the LR system. In this study, we refer to the former (i.e. we do not change the PG software models themselves). However, in the process of creating some of the metrics to assess this, LR-optimisation is employed.

An important aspect is that calibration can only be evaluated when there are enough data points. For example, to ascertain that the proportion of LRs equal to a 10^6^ under H_d_ is around 1/10^6^, we need around a million H_d_-true LRs. Fiducial calibration discrepancy plots and PAV plots make this data dependence very explicit by only showing performance for the log_10_LR ranges in which LR values exist for both H_p_-true and H_d_-true scenarios [30, 31, 32]. Beyond these limits, there are insufficient data to evaluate calibration performance [31]. In theory, ECE plots and calibration tables indicate calibration across the full range of LRs [27, 29], yet these metrics are still limited by dataset size.

## 2. Materials and methods

### 2.1 DNA profiles

In this study, LRs were calculated with DNA mixture profiles in the publicly available Boston University PROVEDIt Initiative (Project Research Openness for Validation with Empirical Data) dataset. In many cases the mixtures were deteriorated with treatments (DNAse I degradation, Fragmentase® degradation, UV damage, sonification, or humic acid inhibition) [36].

For each mixture in this dataset, the number of contributors (NoC), sample mixture ratio, and DNA quantification results are specified. As in Susik and Sbalzarini [17], we analysed 428 mixtures that were amplified with the GlobalFiler kit (29 PCR cycles) and analysed on a 3500 Genetic Analyser (15s injection time). We used the same input files as in [17]. For use in DNAStatistX, we removed any empty cell spaces present as these would be incorrectly parsed by the software. The reference profiles of all true donors (N=22) to the PROVEDIt DNA mixture profiles were available, as well as a set of non-contributor DNA profiles (N=715). In addition, random non-contributor DNA reference profiles were simulated using the NIST 1036-Caucasian allele frequencies database. In total, 500 of these simulated reference profiles were used in this study.

### 2.2 LR calculations

LRs were computed using DNAStatistX v2.1.0 and EuroForMix v4.0.8 with hypothesis sets:

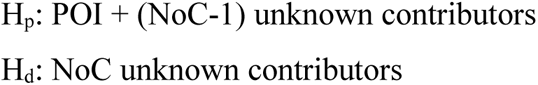

NoC was selected as the “true NoC” (i.e as described by the sample) unless stated otherwise. Settings for LR calculations are presented in Table 1.

**Table 1.** Settings for LR calculations in DNAStatistX v2.1.0 and EuroForMix v4.0.8.

| Setting |  |
| --- | --- |
| Allele frequency file<sup>a</sup> | NIST 1036-Caucasian allele frequencies based on 361 individuals |
| Rare allele frequency<sup>a</sup> | 0.006925208 |
| Fst | 0.01<sup>a</sup> for both software, in addition 0.03 using DNASTatistX |
| Drop-in probability<sup>a</sup> | 0.00073 |
| Drop-in lambda<sup>a</sup> | 0.03846 |
| Kit<sup>a</sup> | GlobalFiler |
| Analytical thresholds (RFU)<sup>a</sup> | D3S1358=60, vWA=60, D16S539=60, CSF1PO=60, TPOX=60, D8S1179=80, D21S11=80, D18S51=80, D2S441=45, D19S433=45, TH01=45, FGA=45, D22S1045=75, D5S818=75, D13S317=75, D7S820=75, SE33=75, D10S1248=100, D1S1656=100, D12S391=100, D2S1338=100 |
| Model | MLE |
| Degradation model | ON for DNASTatistX. ON for EuroForMix, except if degradation slope estimated >0.99, then OFF |
| Stutter models | By model selection, largest adjusted log likelihood out of stutter models OFF, BW, FW, BW & FW |

<sup>a</sup>Setting as in [17].

LR calculations were performed with each mixture for:

(i) each true-contributor reference profile as the POI (H_p_-true scenario); and
(ii) non-contributor reference profiles as the POI (H_d_-true scenario).

With respect to (ii), the number of non-contributor reference profiles for which LRs were calculated was equal to the NoC. I.e. if NoC = 4, a total of eight separate LRs were computed with that profile; one with each of the four true contributors and four with a different non-contributor as the POI.

#### 2.2.1 DNAStatistX and EuroForMix model selection and model validation

Model selection is an exploratory step carried out during an LR calculation, to determine the combination of model settings that best explain the input data. In this study, model selection was performed using DNAStatistX and EuroForMix independently. DNAStatistX model selection was performed according to casework practice for DNA profiles where no stutter filtering was applied during analysis. For efficiency of calculations, EuroForMix model selection took a slightly different approach, using all true contributors to reach the optimum model conditions (discussed in detail below). As a result, DNAStatistX and EuroForMix LRs were calculated using – for the majority of hypotheses – the same selected models, yet not always. These different approaches may cause variation in LRs, though similar trends are expected.

Model validation, on the other hand, is an aspect of DNAStatistX and EuroForMix software which is performed on a per case basis, and plots the cumulative probabilities for the expected peak heights against those for the observed peak heights. A Bonferroni corrected significance level of 0.01 is applied, and if four or more data points fall outside this envelope, the model validation is scored as failed.

For model selection in DNAStatistX, the degradation model was turned ON in all computations and Fst (‘θ’) set to 0.01, whilst the stutter models (backward (BW) and/or forward (FW)) were turned ON and OFF to infer which model combination enabled the best explanation of the data (i.e. no stutter models (SM OFF), BW, FW, BW & FW). This yielded a total of 10,056 LRs (2,514 as in [17] times four model conditions) (Table 2). For each scenario (i.e. set of propositions for mixture + POI), the model condition that yielded the largest adjusted log likelihood under H_d_ and model validation passed was selected (similar to [13]). For the rest of the scenarios that failed model validation, the best model condition was selected (largest adjusted log likelihood under H_d_), unless it was an H_p_-true scenario, or an H_d_-true scenario with a log_10_LR >-4.

**Table 2.**
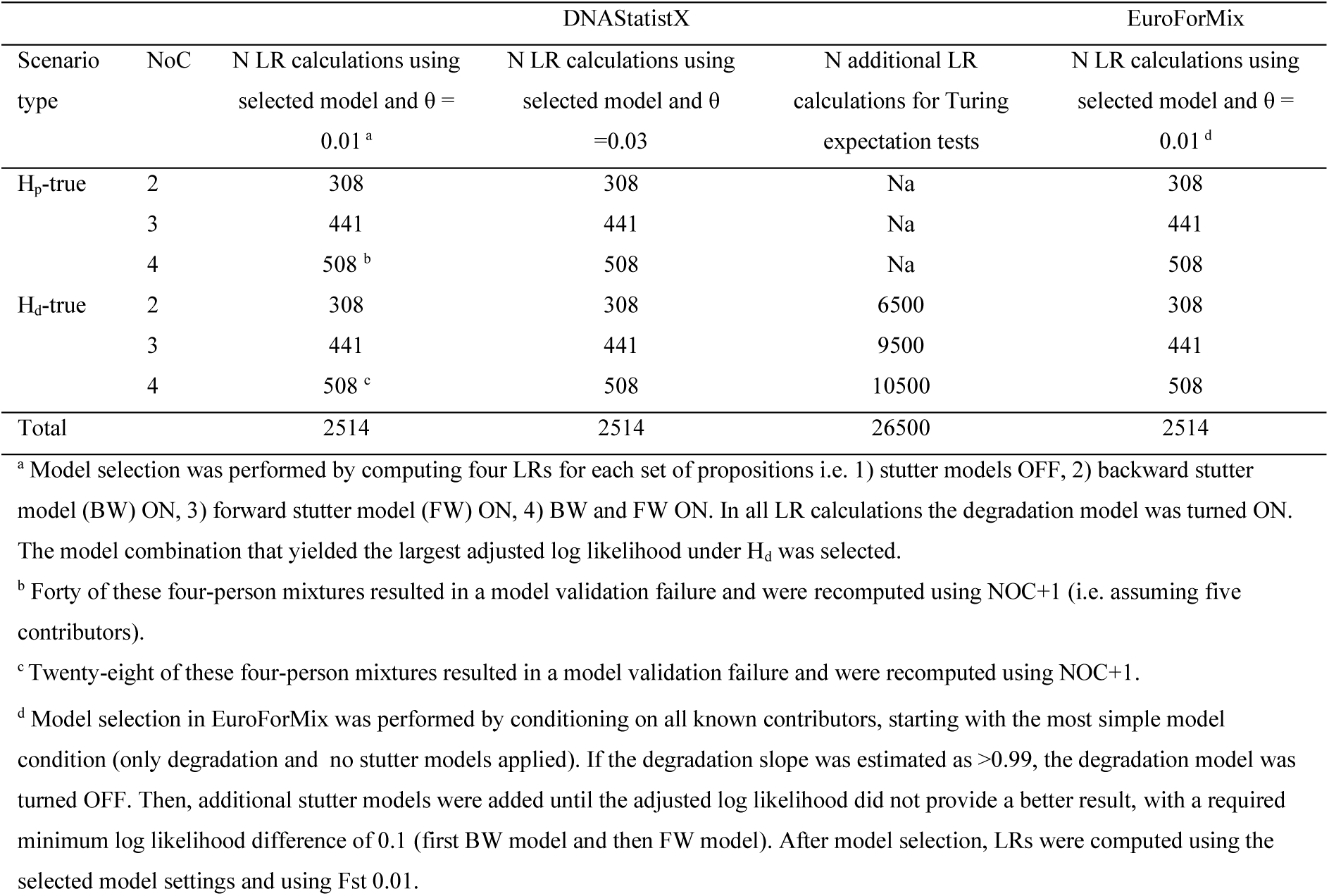
Number of LR calculations performed of each scenario type.

For H_p_-true LR scenarios with a failed model validation and H_d_-true scenarios with log_10_LR>-4 (N=17) additional LR calculations were performed using NoC+1 for each model condition. This yielded a total of four additional LR calculations per failed model validation result (N=68). If model validation passed under any NoC+1 model condition, the best model condition of these passes (largest log likelihood under H_d_) was selected. If none of the NoC+1 calculations passed model validation, then the model condition with just the largest log likelihood under H_d_ was selected out of the original LRs and the NoC+1 scenarios (this process is summarised in Supplementary Fig. 1).

In casework practice, it is advised to further examine LR results that are accompanied with a failed model validation and to not report the LR if the failure cannot be solved or explained. In this study, we kept all results enabling comparison to previously reported results using other models. Furthermore, we examined whether observed trends differed if the data with failed model validations were left out.

Lastly, DNAStatistX LR calculations were repeated with the final selected model conditions and using a larger value for Fst (θ = 0.03 instead of θ = 0.01). Different Fst (θ) values may be employed depending on the relevant population database for a case as well as the results of internal validation studies [17]. At the authors’ institute using DNAStatistX, θ = 0.03 is applied in casework, and in the article of Susik & Szbalzarini θ = 0.01 was used. Ultimately, this created two conditions: ‘DNAStatistX θ = 0.01’ and ‘DNAStatistX θ = 0.03’.

Susik and Sbalzarini performed EuroForMix v3.4 calculations [17]. However, in the current study, calculations were rerun in EuroForMix v4.0.8 (hereafter referred to as EuroForMix) with a slightly different strategy for model selection (Table 2). Susik and Sbalzarini started with a saturated model (always using the degradation model and both stutter models) and reduced the model conditions until it converged. They did not perform model selection for the four-person mixtures and for none of the LRs, the model validation results were reported [17]. In the current study, the model selection for EuroForMix was carried out by conditioning on the true contributors and increasing the complexity if the log-likelihood was increasing with a value of 0.01. Additionally, the degradation model was turned off if the estimated degradation slope was above 0.99. These strategies were applied to avoid convergence issues. This strategy was performed for all mixture profiles. And for all scenarios, the model validation results were evaluated.

Information on model selection and model validation are presented in Supplementary Table 1. Data for comparisons with HMC and STRmix came from supplementary data and table 1 in Susik and Sbalzarini [17].

### 2.3 Discriminatory power

For both the DNAStatistX and EuroForMix LR results, the rate of misleading evidence was calculated and compared to results from Susik and Sbalzarini [17]. Misleading evidence are also denoted as false positives (FP) and false negatives (FN), or as Type II and Type I errors, respectively [1, 14]. In [17], authors denoted the rate of misleading evidence the “Opposite of the Neutral Threshold” (OotNT). The neutral threshold is LR = 1 (log_10_LR = 0). Misleading evidence (or OotNT scenarios) occurs when the software outputs an LR > 1 (log_10_LR > 0) for an H_d_-true scenario (false positive) or LR < 1 (log_10_LR < 0) for an H_p_-true scenario (false negative). H_d_-true and H_p_-true scenarios that were misleading were summed separately. The rates of misleading evidence were calculated as the proportion of all scenarios that yielded misleading evidence. When discrimination is perfect, this rate will be 0.

Sensitivity, specificity, PPV, NPV and accuracy were calculated as:

- Sensitivity = true positives / (true positives + false negatives)
- Specificity = true negatives / (false positives + true negatives)
- PPV = true positives/(true positives + false positives)
- NPV = true negatives/(true negatives + false negatives)
- Accuracy = (true positives + true negatives) / (true positives + true negatives + false positives + false negatives)

### 2.4 Calibration metrics

Calibration performance of the PG software was assessed by generating calibration tables, fiducial calibration discrepancy plots, ECE plots (with Cllr and Cllr*^cal^* values), PAV plots (with devPAV values) and Turing tests as described in section 1.2.5. Turing tests were only performed for DNAStatistX, however, due to computational load.

For comparison between different models, calibration tables and plots were generated per PG model using the data in [17], for HMC and STRmix, as well as for EuroForMix and each DNAStatistX condition (θ =0.01, θ = 0.03).

Since calibration performance can be dependent on the dataset size (see section 1.2.6), we employed extended DNAStatistX datasets for these metrics.

#### 2.4.1 Interpretation of calibration tables

The observed relative frequencies of H_p_-true scenarios were scored as ‘correct’, ‘more’ and ‘fewer’ (as in [37]) which means that the observed frequency was within, higher than, and less than the range of the corresponding posterior probability range, respectively. Ideally, all observed relative frequencies are within the corresponding posterior probability range, following mathematical expectations (i.e. are ‘correct’).

In this study, we added a further interpretation strategy. The calibration table results were further interpreted as ‘expected’, ‘conservative’, ‘anti-conservative’, and ‘not enough data’. Results were ‘expected’ in posterior probability ranges where ‘correct’ scores were given. If the score was ‘more’ or ‘fewer’ in log_10_LR ranges >0 and <0 respectively, we interpreted this as ‘conservative’. That is, fewer counts of misleading evidence were observed than expected, given the number of H_p_-true counts in that posterior probability range. Importantly, in this context, ‘conservative’ results were more favorable than ‘anti-conservative’.

However, ‘correct’, ‘conservative’, and ‘anti-conservative’ interpretations can only be made if the data size is adequate to assess calibration in a given posterior probability range. If a ‘more’ score changed to ‘fewer’ by adding one additional count to the H_d_-true scenarios in log_10_LR ranges >0 (and vice versa in log_10_LR ranges <0), we assigned ‘not enough data’. Note that in more data-driven forensic fields, the upper threshold for LR reporting is found using this concept of adding one misleading data point – LR ranges denoted ‘not enough data’ here would be outside of the range suitable to report [38].

We further note that the posterior probability range encompassing log_10_LR 0, as well as slightly above and below, were labelled ‘NA’ (not applicable) as these could not be given the interpretation conservative or anti-conservative.

## 3. Results

### 3.1 Discriminatory power analysis of PG models

#### 3.1.1 Comparison of PG models

In this study, DNAStatistX and EuroForMix (EFM) LRs were computed using the data and scenarios as described in [17] and compared to the results that Susik and Sbalzarini previously presented for HMC, and STRmix. Histograms of log_10_LRs from each PG software are shown in Supplementary Fig. 2. In the HMC and STRmix results, log_10_LRs reported as –∞ were set to –200 as the low maximum.

Fig. 4 compares DNAStatistX LRs to those of other PG models at θ = 0.01. Variation in LRs is observed, though LRs for the H_p_-true scenarios are more similar between models than H_d_-true scenario LRs. Specifically, the H_d_-true LRs are higher in DNAStatistX and EuroForMix than in HMC and STRmix. As expected, overall, the DNAStatistX results are most similar to EFM results (Fig. 4). In the next section, we focus on comparing the false positive and false negative results.

**Figure 4.**
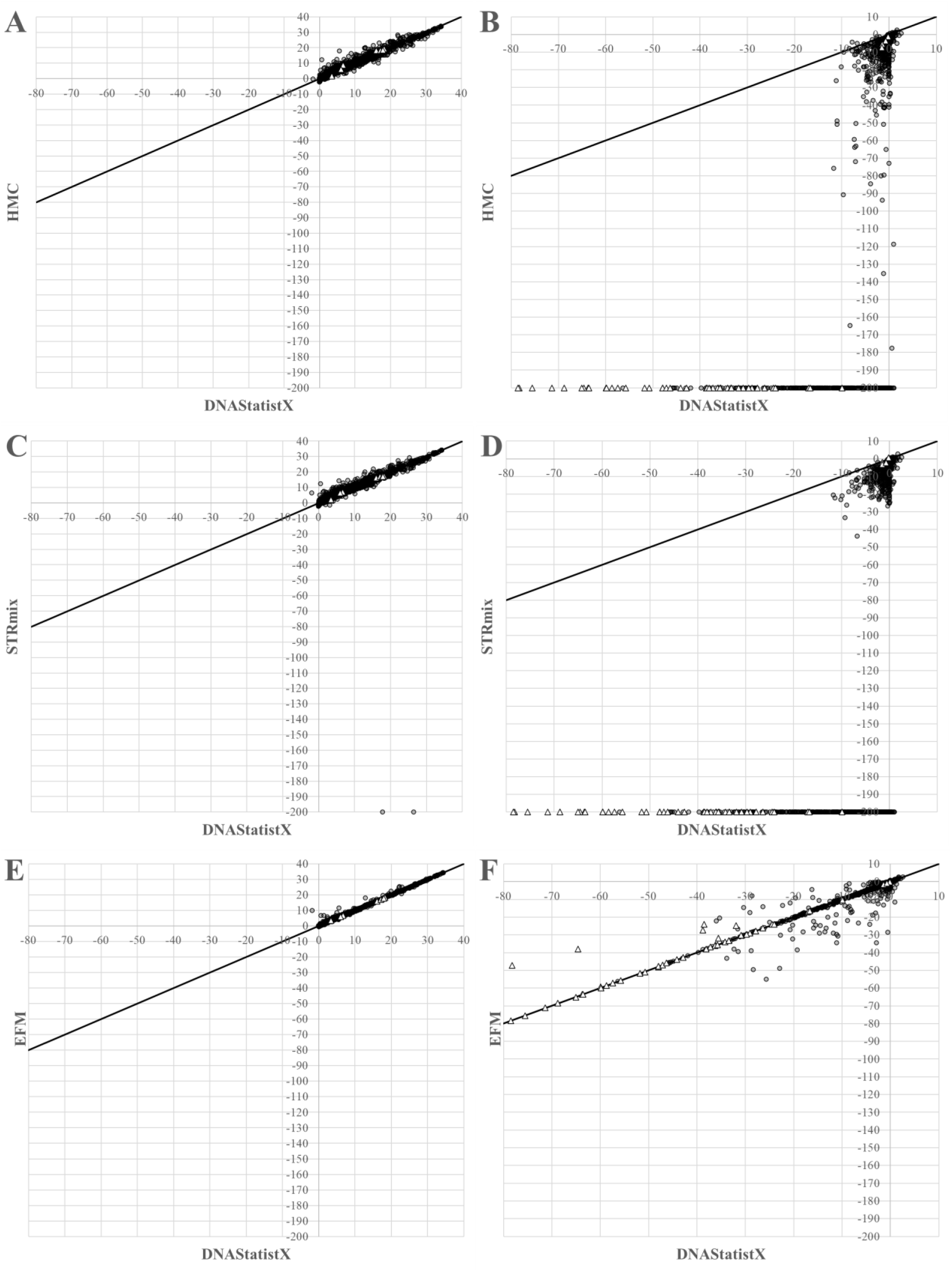
Scatterplots comparing DNAStatistX θ = 0.01 log_10_LRs (X-axes) to results obtained using EuroForMix (EFM) in this study and the log_10_LRs reported for HMC and STRmix as presented in [17] (Y-axes). Results for H_p_-true tests (A, C, E) and H_d_-true tests (B, D, F). Passed and failed model validation in DNAStatistX is presented as circles and triangles, respectively. Black diagonal line: X=Y line.

#### 3.1.2 False positives and false negatives

In this section, rates for misleading evidence were computed as in [17] and compared between PG software results.

For all models, as expected, the discrimination was less good in lower LR ranges and when profiles have higher NoCs. In the lower LR ranges, rates for misleading evidence were lowest for HMC and STRmix, followed by DNAStatistX and EuroForMix (Table 3). When log_10_LRs were ≥3, all PG software indicated perfect discrimination, with misleading evidence rates of 0 (no false positives).

**Table 3.**
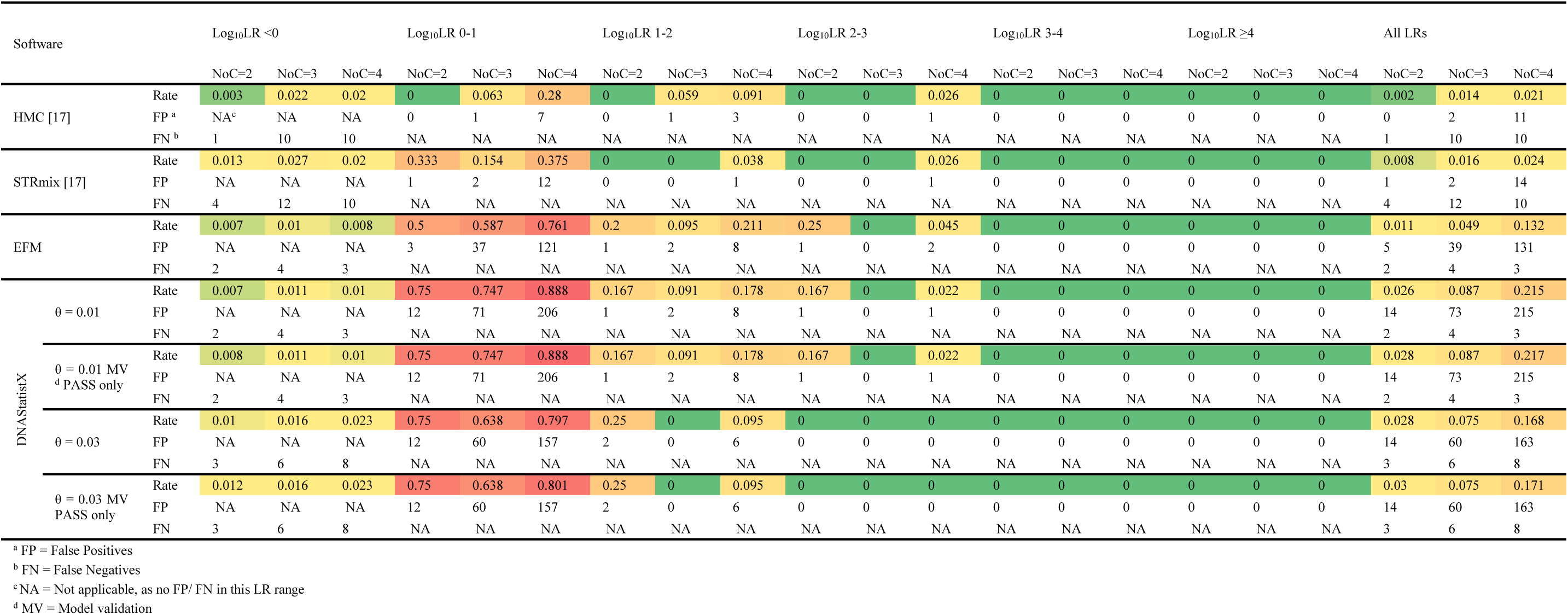
Misleading evidence rates and raw counts for given LR ranges. Perfect discrimination has rate 0, presented in dark green. Lowest to highest rates are shown on a scale from dark green, to light green, to yellow, to orange, to red.

In DNAStatistX and EuroForMix results, false positives were much more common than false negatives. Furthermore, more false positives were observed in DNAStatistX and EuroForMix than other software (Table 3). However, the LR values of these false positives, were lower for DNAStatistX than HMC and STRmix, though DNAStatistX and EuroForMix showed more outliers (Fig. 5). In DNAStatistX, false positive log_10_LRs (θ = 0.01) were at maximum 2.16, 1.26 and 2.57, for two-, three-, and four-person mixtures, respectively. This is within the same range as values that were previously reported for EuroForMix using θ = 0.01 (1.81 and 2.08 for three– and four-person mixtures, respectively [14]). However, unlike in [14], we did observe false positives for two-person mixtures. On the other hand, numbers of false negatives were similar across all software, and sometimes lower in DNAStatistX and EuroForMix (Table 3).

**Figure 5.**
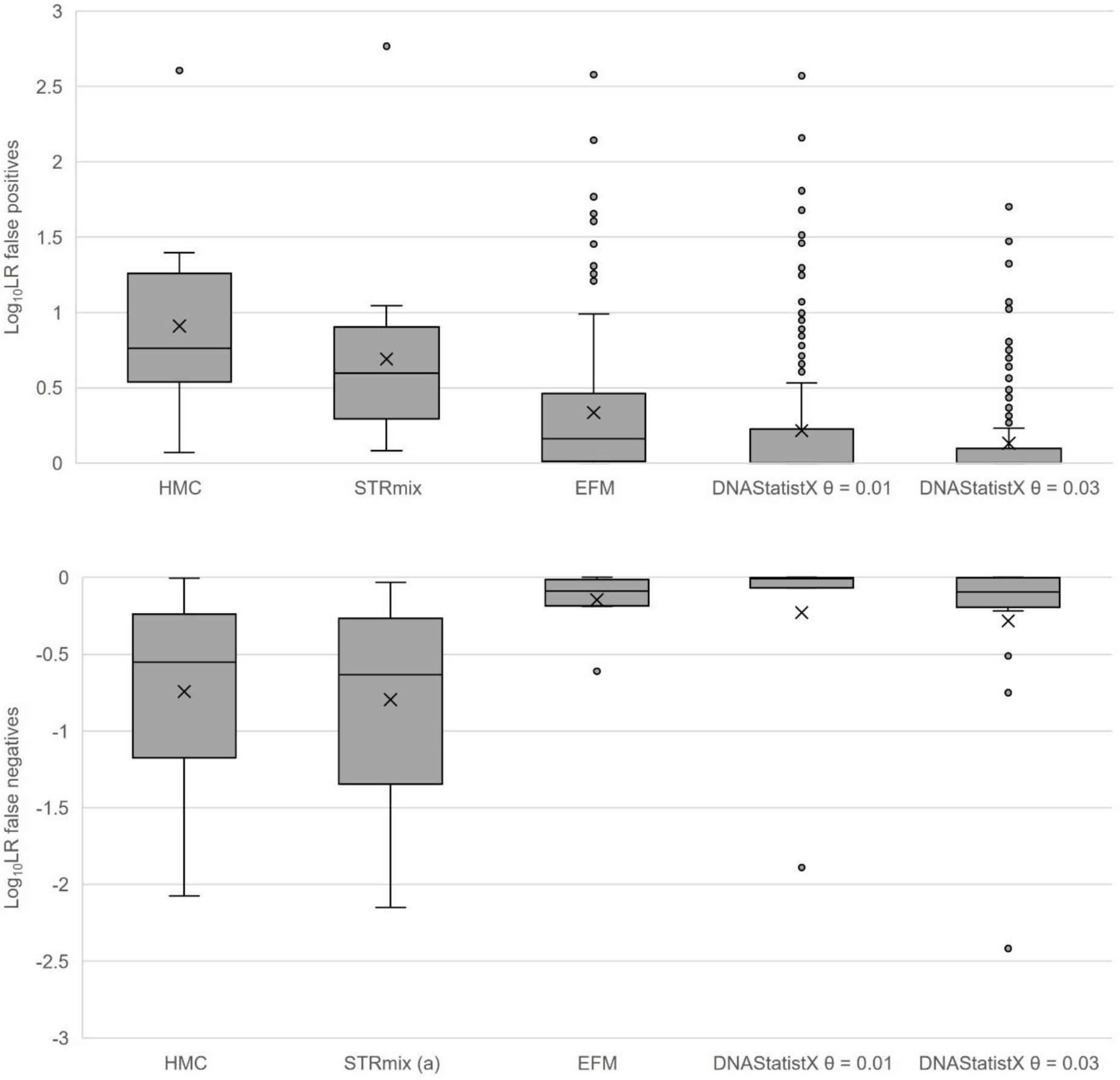
False positive (A) and false negative (B) log_10_LR ranges under each software. (a) _-∞_ in 2 scenarios were omitted from the graph for readability.

In DNAStatistX, as expected, as LRs decreased and with a larger value for Fst, the numbers of false positives decreased and false negatives increased. Rates for misleading evidence were generally lower when using θ = 0.03 than θ = 0.01, especially for three– and four-person mixtures. Furthermore, there were no false positives with log_10_LRs ≥2 when using θ = 0.03, and none with log_10_LRs ≥3 when using θ = 0.01 (Table 3 and Fig. 6). When considering all LRs, DNAStatistX and EuroForMix had higher scores for sensitivity and negative predictive value (NPV) with three– and four-person mixtures, though lower scores for specificity, positive predictive value (PPV) and accuracy (Supplementary Table 2). Trends with respect to DNAStatistX were independent of the few failed model validation results (Table 3), therefore we performed analyses beyond Table 3, with all LRs included (i.e. passes and fails).

**Figure 6.**
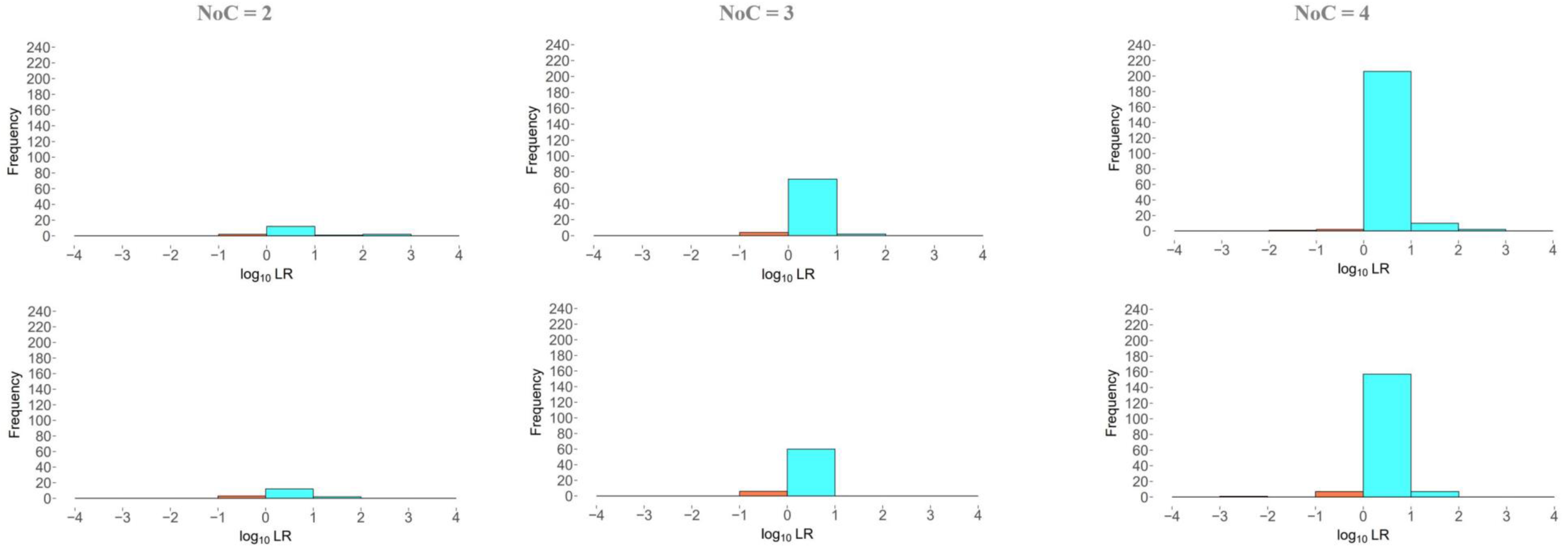
Histograms of log_10_LR results of DNAStatistX false positives (cyan; H_d_-true scenarios with LR >1) and false negatives (orange; H_p_-true scenarios with LR <1). Top: θ = 0.01, Bottom: θ = 0.03.

Consistent with rates for misleading evidence (Table 3), based on the AUCs in Fig. 7, discrimination performance was closer to perfect performance for HMC and STRmix, closely followed by EuroForMix and DNAStatistX.

**Figure 7.**
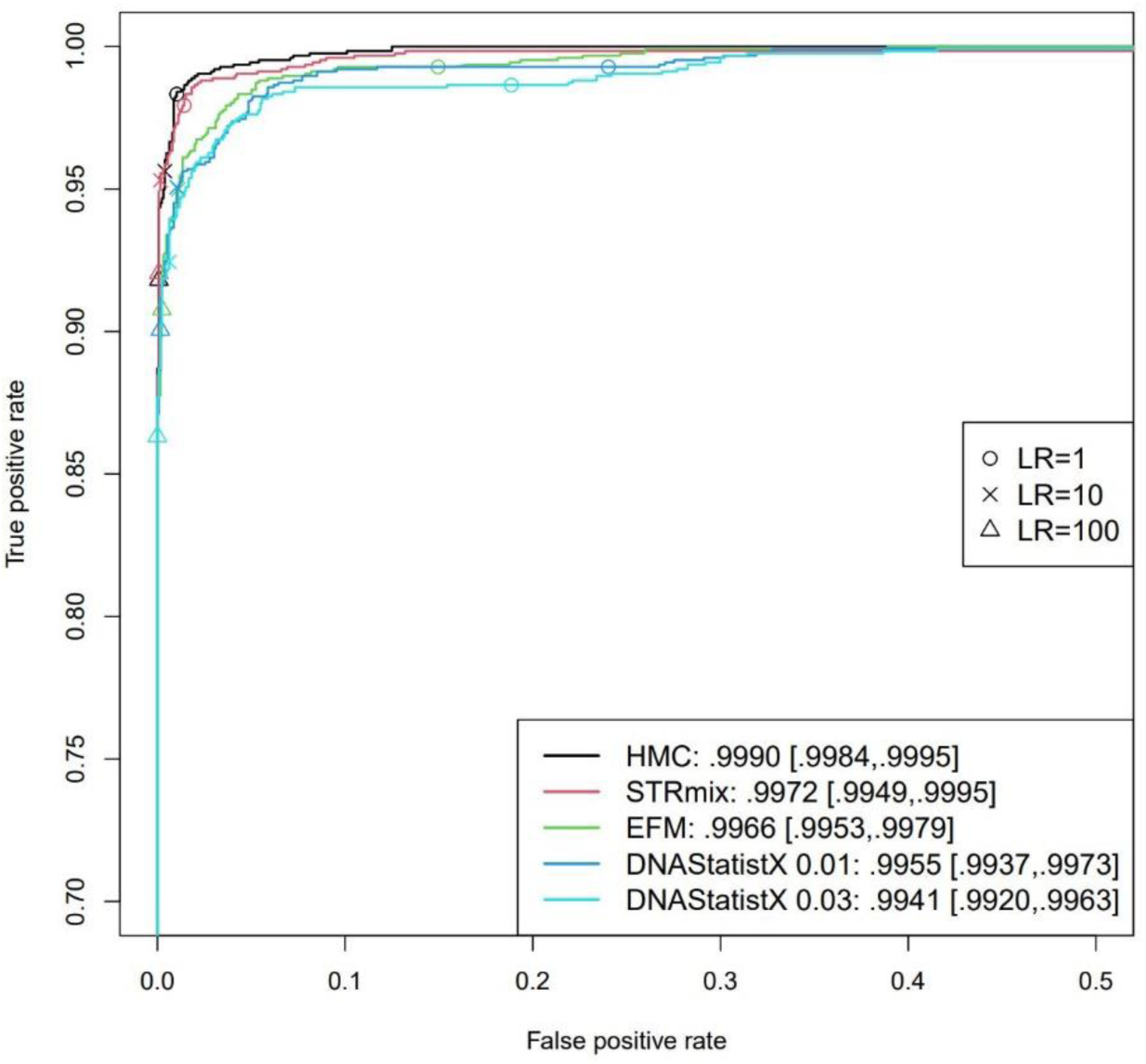
ROC curve of PG models. The numbers in the legend show the estimated AUC with a 95% confidence interval in bracket.

Before continuing, we note an important decision regarding the two H_p_-true scenarios that yielded LRs of –∞ in STRmix. Based on manual comparison of the trace and reference profiles of these scenarios and the results obtained by the other software systems, these results seem unlikely and may be incorrect. In fact, in [37], the results for these same two profile were amended as the authors believed this was closer to correct analysis. Importantly, calibration and ROC plots in the current study are negatively affected by leaving these two datapoints in. Therefore, we decided to omit these very unexpected LR results from these profiles beyond Fig. 6.

### 3.2 Calibration analysis based on calibration tables

#### 3.2.1 Calibration performance using DNAStatistX

To assess the calibration performance of DNAStatistX, we start with calibration tables as presented in [29]. We used the scenarios as described in [17] with the addition of H_d_-true scenarios from Turing expectation non-contributor tests to increase the data size. We refer to this as the ‘extended’ dataset. As expected given the dataset size limitations, all observed relative frequencies of H_p_-true donors in the range log_10_LR ≥3 were 1 and interpreted as ‘not enough data’. The same holds for the lower posterior probability ranges; whereby all observed relative frequencies of H_p_-true donors in the range log_10_LR < –3 were 0, with this same interpretation (Table 4A). Therefore, results show that the dataset size was too limited to assess calibration in the extreme high and low ranges of LRs. In theory, to obtain an LR of e.g. a million for an H_d_-true scenario, at least a million H_d_-true tests should be performed. In practice, such large numbers of tests is not doable for the current PG systems due to computational requirements. In line with this idea, Bright at al. [29] reported no calibration-issues with posterior probabilities above 0.99, resulting in observed relative frequency of H_p_-true donors of 1. With this in mind, and until computational load is further decreased, we note that assessing calibration performance of PG systems is only doable for ‘mid-range’ LRs from ∼log_10_ –3 to log_10_ +3.

**Table 4.** Counts and observed relative frequency of H_p_-true donor scenarios and the count of H_d_-true donor scenarios with posterior probability results Pr (Hp∣LR) within each range when LRs were computed using the extended DNAStatistX datasets. A) θ = 0.01, B) θ = 0.03.

| **A** DNASTatistX $\theta = 0.01$ | | | | | | | |
| --- | --- | --- | --- | --- | --- | --- | --- |
| Posterior probability range $\text{Pr}(H_p\vert \text{LR})$ | $\log_{10}\text{LR}$ interval (2dp) | LR interval | $H_d$-true donors Count | $H_p$-true donors Count | Observed relative frequency of $H_p$-true donors | Comment regarding $H_p$-true donors | Interpretation |
| 0.999999 to 1 | 7.02 to ∞ | 10546529 to ∞ | 0 | 781 | 1 | Correct | Not enough data <sup>b</sup> |
| 0.99999 to 0.999999 | 6.02 to 7.02 | 1054643 to 10546529 | 0 | 67 | 1 | More | Not enough data |
| 0.9999 to 0.99999 | 5.02 to 6.02 | 105455 to 1054643 | 0 | 81 | 1 | More | Not enough data |
| 0.999 to 0.9999 | 4.02 to 5.02 | 10536 to 105455 | 0 | 74 | 1 | More | Not enough data |
| 0.99 to 0.999 | 3.02 to 4.02 | 1044 to 10536 | 0 | 71 | 1 | More | Not enough data |
| 0.95 to 0.99 | 2.3 to 3.02 | 200 4 to 1044 | 3 | 37 | 0.93 | Fewer | Anti-conservative <sup>c</sup> |
| 0.75 to 0.95 | 1.5 to 2.3 | 31.6 to 200 | 15 | 51 | 0.77 | Correct | Expected <sup>d</sup> |
| 0.25 to 0.75 | 0.55 to 1.5 | 3.5 to 31.6 | 253 | 51 | 0.17 | Fewer | Anti-conservative |
| 0.05 to 0.25 | -0.26 to 0.55 | 0.6 to 3.5 <sup>a</sup> | 7997 | 43 | 0.0050 | Fewer | N/A |
| 0.01 to 0.05 | -0.97 to -0.26 | 0.1 to 0.6 | 954 | 0 | 0 | Fewer | Conservative <sup>e</sup> |
| 0.001 to 0.01 | -1.98 to -0.97 | 0.01 to 0.1 | 673 | 1 | 0.0015 | Correct | Expected |
| $1.00 \times 10^{-4}$ to $1.00 \times 10^{-3}$ | -2.98 to -1.98 | 0.001 to 0.01 | 705 | 0 | 0 | Fewer | Not enough data |
| $1.00 \times 10^{-5}$ to $1.00 \times 10^{-4}$ | -3.98 to -2.98 | 0.0001 to 0.001 | 781 | 0 | 0 | Fewer | Not enough data |
| $1.00 \times 10^{-7}$ to $1.00 \times 10^{-5}$ | -5.98 to -3.98 | 0.000001 to 0.00010 | 957 | 0 | 0 | Fewer | Not enough data |
| 0 to $1.00 \times 10^{-7}$ | $-\infty$ to -5.98 | $-\infty$ to 0.000001 | 419 | 0 | 0 | Fewer | Not enough data |
| **B** DNASTatistX $\theta = 0.03$ | | | | | | | |
| Posterior probability range $\text{Pr}(H_p\vert \text{LR})$ | $\log_{10}\text{LR}$ interval (2dp) | LR interval | $H_d$-true donors Count | $H_p$-true donors Count | Observed relative frequency of $H_p$-true donors | Comment regarding $H_p$-true donors | Interpretation |
| 0.999999 to 1 | 7.12 to ∞ | 13330933 to ∞ | 0 | 676 | 1 | Correct | Not enough data |
| 0.99999 to 0.999999 | 6.12 to 7.12 | 1333081 to 13330933 | 0 | 67 | 1 | More | Not enough data |
| 0.9999 to 0.99999 | 5.12 to 6.12 | 133296 to 1333081 | 0 | 86 | 1 | More | Not enough data |
| 0.999 to 0.9999 | 4.12 to 5.12 | 13318 to 133296 | 0 | 85 | 1 | More | Not enough data |
| 0.99 to 0.999 | 3.12 to 4.12 | 1320 to 13318 | 0 | 90 | 1 | More | Not enough data |
| 0.95 to 0.99 | 2.4 to 3.12 | 253 to 1320 | 0 | 61 | 1 | More | Conservative |
| 0.75 to 0.95 | 1.6 to 2.4 | 40 to 253 | 5 | 45 | 0.90 | Correct | Expected |
| 0.25 to 0.75 | 0.65 to 1.6 | 4.4 to 40 | 115 | 80 | 0.41 | Correct | Expected |
| 0.05 to 0.25 | -0.15 to 0.65 | 0.7 to 4.4 | 7650 | 62 | 0.0080 | Fewer | N/A |
| 0.01 to 0.05 | -0.87 to -0.15 | 0.1 to 0.7 | 3719 | 4 | 0.0010 | Fewer | Conservative |
| 0.001 to 0.01 | -1.87 to -0.87 | 0.01 to 0.1 | 2125 | 0 | 0 | Fewer | Conservative |
| $1.00 \times 10^{-4}$ to $1.00 \times 10^{-3}$ | -2.88 to -1.87 | 0.001 to 0.01 | 723 | 1 | 0.0014 | More | Anti-conservative |
| $1.00 \times 10^{-5}$ to $1.00 \times 10^{-4}$ | -3.88 to -2.88 | 0.0001 to 0.001 | 665 | 0 | 0 | Fewer | Not enough data |
| $1.00 \times 10^{-7}$ to $1.00 \times 10^{-5}$ | -5.88 to -3.88 | 0.000001 to 0.00010 | 433 | 0 | 0 | Fewer | Not enough data |
| 0 to $1.00 \times 10^{-7}$ | $-\infty$ to -5.88 | $-\infty$ to 0.000001 | 822 | 0 | 0 | Fewer | Not enough data |

<sup>a</sup> Grey text = LRs below $\log_{10} 0$ (approximate due to rounding).
<sup>b</sup> Not enough data = in theory, more $H_p$ -true counts in this range are needed to observe an $H_d$ -true result/ more $H_p$ -true counts in this range are needed to be able to observe an $H_p$ -true result.
<sup>c</sup> Anti-conservative = more false-positives observed than expected based on counts of $H_p$ -true donors in this range/ more false negatives were observed than expected based on the counts of $H_p$ -true donors in this range.
<sup>d</sup> Expected = The observed relative frequency of $H_p$ -true donors falls within the posterior probability range.
<sup>e</sup> Conservative (too close to LR=1) = with the number of $H_p$ -true counts in this range a false-positive would be expected but was not observed/ with the number of $H_d$ -true counts in this range a false-negative would be expected but was not observed.

Examination of the mid-range LRs in the calibration tables for DNAStatistX shows a better calibration performance with θ = 0.03 rather than θ = 0.01. Firstly, the DNAStatistX θ = 0.03 calibration table shows more ‘correct’ scorings in the log_10_LR range –3 to +3 than the DNAStatistX θ = 0.01 calibration table (Tables 4A and 4B). Furthermore, DNAStatistX θ = 0.01 shows anti-conservative results in the log_10_LR range ∼0-3 (Table 4A). On the other hand, switching to θ = 0.03 eliminated all anti-conservative results for log_10_LR >0 (Table 4B). Although θ is a subpopulation correction, these results indicate that it can be effective as a correction factor for ‘low’ LRs obtained using DNAStatistX.

#### 3.2.2 Comparison of calibration tables for different LR models

To enable comparison between models, the original dataset described by [17] was used. For all software, as expected – given the aforementioned data limitations – relative frequencies of H_p_-true donors were observed to be 1 in the range log_10_LR ≥3, and 0 in the range log_10_LR < –3 (Table 5). Therefore, as concluded in the previous section, calibration tables as presented in [29] and using PG systems’ data seem to be most useful to assess mid-range LRs (∼log_10_ –3 to +3).

**Table 5.**
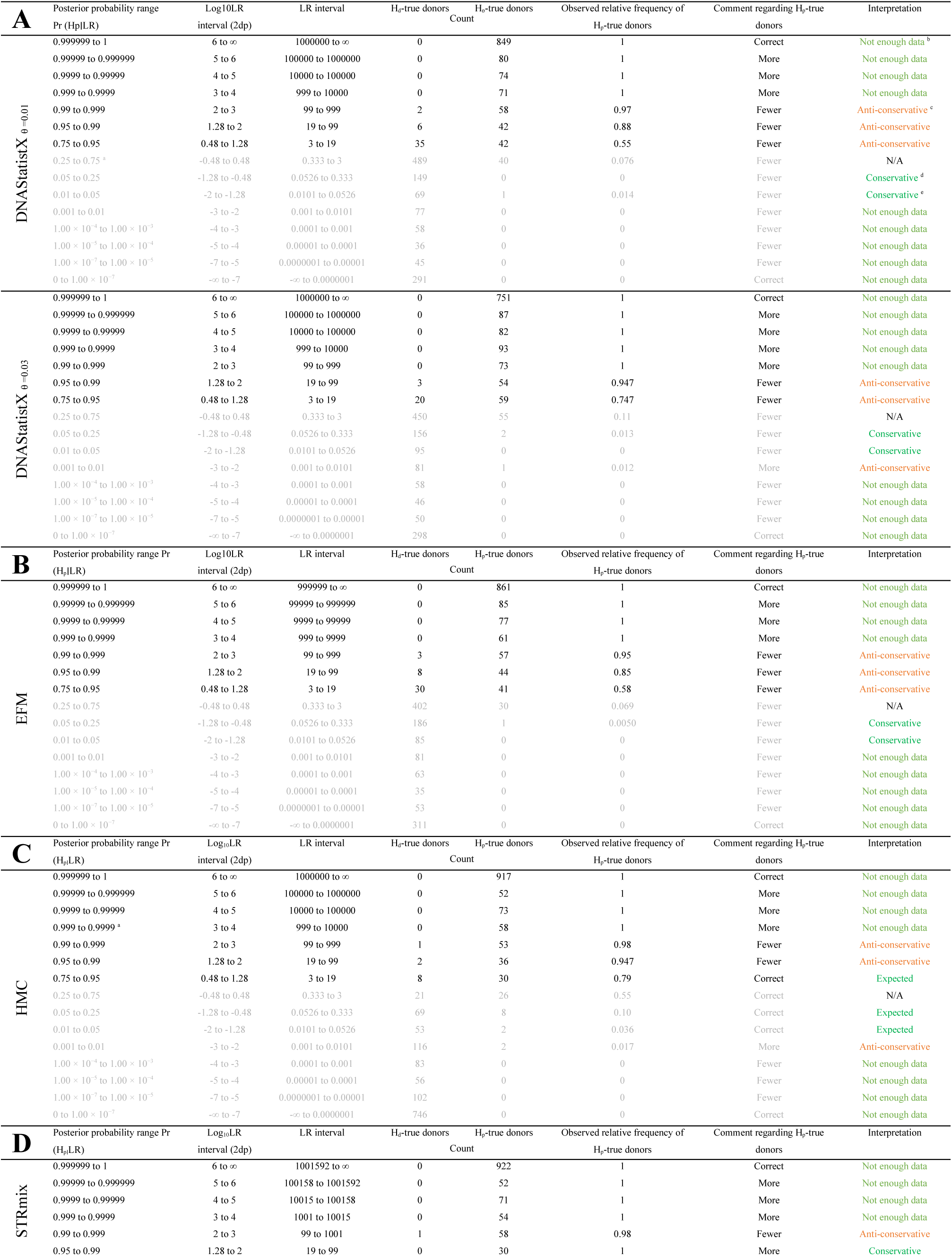
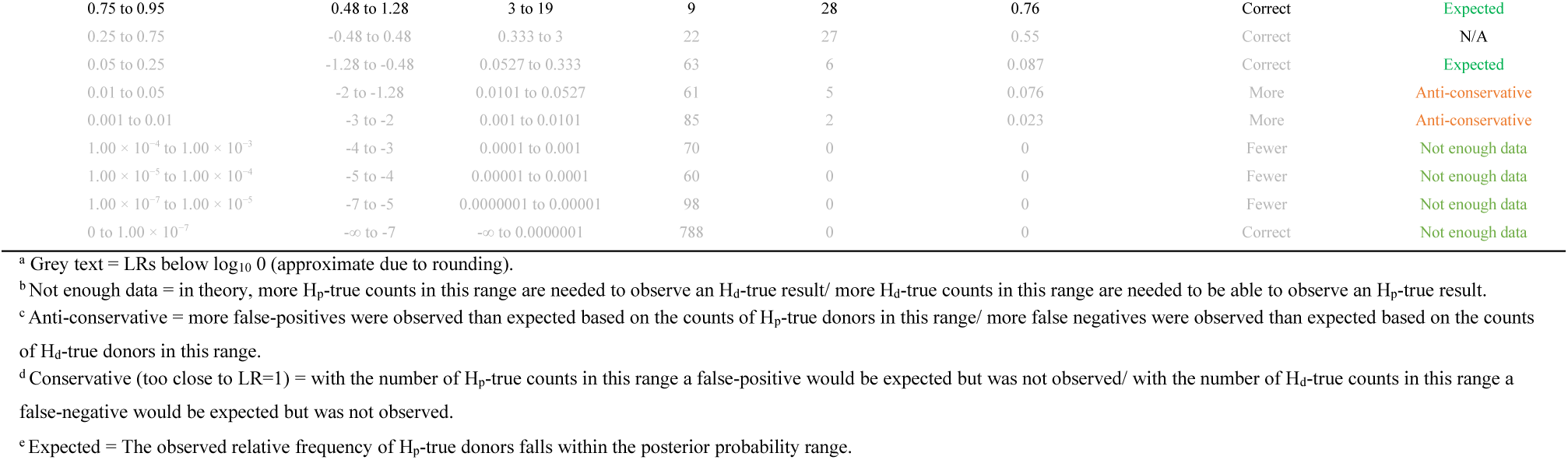
Counts and observed relative frequency of H_p_-true donor scenarios and the count of H_d_-true donor scenarios with posterior probability results Pr(H_p_∣LR) within each range when LRs were computed using A) DNAStatistX (θ =0.01 & θ =0.03), B) EuroForMix, C) HMC [17], D) STRmix [17].

| **A** | Posterior probability range<br>Pr ($H_p\vert LR$) | Log10LR<br>interval (2dp) | LR interval | $H_d$-true donors<br>Count | $H_p$-true donors<br>Count | Observed relative frequency of<br>$H_p$-true donors | Comment regarding $H_p$-true<br>donors | Interpretation |
| --- | --- | --- | --- | --- | --- | --- | --- | --- |
| DNASStatistX $\theta=0.01$ | 0.999999 to 1 | 6 to ∞ | 1000000 to ∞ | 0 | 849 | 1 | Correct | Not enough data <sup>b</sup> |
|  | 0.99999 to 0.999999 | 5 to 6 | 100000 to 1000000 | 0 | 80 | 1 | More | Not enough data |
|  | 0.9999 to 0.99999 | 4 to 5 | 10000 to 100000 | 0 | 74 | 1 | More | Not enough data |
|  | 0.999 to 0.9999 | 3 to 4 | 999 to 10000 | 0 | 71 | 1 | More | Not enough data |
|  | 0.99 to 0.999 | 2 to 3 | 99 to 999 | 2 | 58 | 0.97 | Fewer | Anti-conservative <sup>c</sup> |
|  | 0.95 to 0.99 | 1.28 to 2 | 19 to 99 | 6 | 42 | 0.88 | Fewer | Anti-conservative |
|  | 0.75 to 0.95 | 0.48 to 1.28 | 3 to 19 | 35 | 42 | 0.55 | Fewer | Anti-conservative |
|  | 0.25 to 0.75 <sup>a</sup> | -0.48 to 0.48 | 0.333 to 3 | 489 | 40 | 0.076 | Fewer | N/A |
|  | 0.05 to 0.25 | -1.28 to -0.48 | 0.0526 to 0.333 | 149 | 0 | 0 | Fewer | Conservative <sup>d</sup> |
|  | 0.01 to 0.05 | -2 to -1.28 | 0.0101 to 0.0526 | 69 | 1 | 0.014 | Fewer | Conservative <sup>e</sup> |
|  | 0.001 to 0.01 | -3 to -2 | 0.001 to 0.0101 | 77 | 0 | 0 | Fewer | Not enough data |
| | $1.00 \times 10^{-4}$ to $1.00 \times 10^{-3}$ | -4 to -3 | 0.0001 to 0.001 | 58 | 0 | 0 | Fewer | Not enough data |
| | $1.00 \times 10^{-5}$ to $1.00 \times 10^{-4}$ | -5 to -4 | 0.00001 to 0.0001 | 36 | 0 | 0 | Fewer | Not enough data |
| | $1.00 \times 10^{-7}$ to $1.00 \times 10^{-5}$ | -7 to -5 | 0.0000001 to 0.00001 | 45 | 0 | 0 | Fewer | Not enough data |
| | 0 to $1.00 \times 10^{-7}$ | $-\infty$ to -7 | $-\infty$ to 0.0000001 | 291 | 0 | 0 | Correct | Not enough data |
| DNASStatistX $\theta=0.03$ | 0.999999 to 1 | 6 to ∞ | 1000000 to ∞ | 0 | 751 | 1 | Correct | Not enough data |
|  | 0.99999 to 0.999999 | 5 to 6 | 100000 to 1000000 | 0 | 87 | 1 | More | Not enough data |
|  | 0.9999 to 0.99999 | 4 to 5 | 10000 to 100000 | 0 | 82 | 1 | More | Not enough data |
|  | 0.999 to 0.9999 | 3 to 4 | 999 to 10000 | 0 | 93 | 1 | More | Not enough data |
|  | 0.99 to 0.999 | 2 to 3 | 99 to 999 | 0 | 73 | 1 | More | Not enough data |
|  | 0.95 to 0.99 | 1.28 to 2 | 19 to 99 | 3 | 54 | 0.947 | Fewer | Anti-conservative |
|  | 0.75 to 0.95 | 0.48 to 1.28 | 3 to 19 | 20 | 59 | 0.747 | Fewer | Anti-conservative |
|  | 0.25 to 0.75 | -0.48 to 0.48 | 0.333 to 3 | 450 | 55 | 0.11 | Fewer | N/A |
|  | 0.05 to 0.25 | -1.28 to -0.48 | 0.0526 to 0.333 | 156 | 2 | 0.013 | Fewer | Conservative |
|  | 0.01 to 0.05 | -2 to -1.28 | 0.0101 to 0.0526 | 95 | 0 | 0 | Fewer | Conservative |
|  | 0.001 to 0.01 | -3 to -2 | 0.001 to 0.0101 | 81 | 1 | 0.012 | More | Anti-conservative |
| | $1.00 \times 10^{-4}$ to $1.00 \times 10^{-3}$ | -4 to -3 | 0.0001 to 0.001 | 58 | 0 | 0 | Fewer | Not enough data |
| | $1.00 \times 10^{-5}$ to $1.00 \times 10^{-4}$ | -5 to -4 | 0.00001 to 0.0001 | 46 | 0 | 0 | Fewer | Not enough data |
| | $1.00 \times 10^{-7}$ to $1.00 \times 10^{-5}$ | -7 to -5 | 0.0000001 to 0.00001 | 50 | 0 | 0 | Fewer | Not enough data |
| | 0 to $1.00 \times 10^{-7}$ | $-\infty$ to -7 | $-\infty$ to 0.0000001 | 298 | 0 | 0 | Correct | Not enough data |
| **B** | Posterior probability range Pr<br>($H_p\vert LR$) | Log10LR<br>interval (2dp) | LR interval | $H_d$-true donors<br>Count | $H_p$-true donors<br>Count | Observed relative frequency of<br>$H_p$-true donors | Comment regarding $H_p$-true<br>donors | Interpretation |
| EFM | 0.999999 to 1 | 6 to ∞ | 999999 to ∞ | 0 | 861 | 1 | Correct | Not enough data |
|  | 0.99999 to 0.999999 | 5 to 6 | 99999 to 999999 | 0 | 85 | 1 | More | Not enough data |
|  | 0.9999 to 0.99999 | 4 to 5 | 9999 to 99999 | 0 | 77 | 1 | More | Not enough data |
|  | 0.999 to 0.9999 | 3 to 4 | 999 to 9999 | 0 | 61 | 1 | More | Not enough data |
|  | 0.99 to 0.999 | 2 to 3 | 99 to 999 | 3 | 57 | 0.95 | Fewer | Anti-conservative |
|  | 0.95 to 0.99 | 1.28 to 2 | 19 to 99 | 8 | 44 | 0.85 | Fewer | Anti-conservative |
|  | 0.75 to 0.95 | 0.48 to 1.28 | 3 to 19 | 30 | 41 | 0.58 | Fewer | Anti-conservative |
|  | 0.25 to 0.75 | -0.48 to 0.48 | 0.333 to 3 | 402 | 30 | 0.069 | Fewer | N/A |
|  | 0.05 to 0.25 | -1.28 to -0.48 | 0.0526 to 0.333 | 186 | 1 | 0.0050 | Fewer | Conservative |
|  | 0.01 to 0.05 | -2 to -1.28 | 0.0101 to 0.0526 | 85 | 0 | 0 | Fewer | Conservative |
|  | 0.001 to 0.01 | -3 to -2 | 0.001 to 0.0101 | 81 | 0 | 0 | Fewer | Not enough data |
| | $1.00 \times 10^{-4}$ to $1.00 \times 10^{-3}$ | -4 to -3 | 0.0001 to 0.001 | 63 | 0 | 0 | Fewer | Not enough data |
| | $1.00 \times 10^{-5}$ to $1.00 \times 10^{-4}$ | -5 to -4 | 0.00001 to 0.0001 | 35 | 0 | 0 | Fewer | Not enough data |
| | $1.00 \times 10^{-7}$ to $1.00 \times 10^{-5}$ | -7 to -5 | 0.0000001 to 0.00001 | 53 | 0 | 0 | Fewer | Not enough data |
| | 0 to $1.00 \times 10^{-7}$ | $-\infty$ to -7 | $-\infty$ to 0.0000001 | 311 | 0 | 0 | Correct | Not enough data |
| **C** | Posterior probability range Pr<br>($H_p\vert LR$) | Log<sub>10</sub>LR<br>interval (2dp) | LR interval | $H_d$-true donors<br>Count | $H_p$-true donors<br>Count | Observed relative frequency of<br>$H_p$-true donors | Comment regarding $H_p$-true<br>donors | Interpretation |
| HMC | 0.999999 to 1 | 6 to ∞ | 1000000 to ∞ | 0 | 917 | 1 | Correct | Not enough data |
|  | 0.99999 to 0.999999 | 5 to 6 | 100000 to 1000000 | 0 | 52 | 1 | More | Not enough data |
|  | 0.9999 to 0.99999 | 4 to 5 | 10000 to 100000 | 0 | 73 | 1 | More | Not enough data |
|  | 0.999 to 0.9999 <sup>a</sup> | 3 to 4 | 999 to 10000 | 0 | 58 | 1 | More | Not enough data |
|  | 0.99 to 0.999 | 2 to 3 | 99 to 999 | 1 | 53 | 0.98 | Fewer | Anti-conservative |
|  | 0.95 to 0.99 | 1.28 to 2 | 19 to 99 | 2 | 36 | 0.947 | Fewer | Anti-conservative |
|  | 0.75 to 0.95 | 0.48 to 1.28 | 3 to 19 | 8 | 30 | 0.79 | Correct | Expected |
|  | 0.25 to 0.75 | -0.48 to 0.48 | 0.333 to 3 | 21 | 26 | 0.55 | Correct | N/A |
|  | 0.05 to 0.25 | -1.28 to -0.48 | 0.0526 to 0.333 | 69 | 8 | 0.10 | Correct | Expected |
|  | 0.01 to 0.05 | -2 to -1.28 | 0.0101 to 0.0526 | 53 | 2 | 0.036 | Correct | Expected |
|  | 0.001 to 0.01 | -3 to -2 | 0.001 to 0.0101 | 116 | 2 | 0.017 | More | Anti-conservative |
| | $1.00 \times 10^{-4}$ to $1.00 \times 10^{-3}$ | -4 to -3 | 0.0001 to 0.001 | 83 | 0 | 0 | Fewer | Not enough data |
| | $1.00 \times 10^{-5}$ to $1.00 \times 10^{-4}$ | -5 to -4 | 0.00001 to 0.0001 | 56 | 0 | 0 | Fewer | Not enough data |
| | $1.00 \times 10^{-7}$ to $1.00 \times 10^{-5}$ | -7 to -5 | 0.0000001 to 0.00001 | 102 | 0 | 0 | Fewer | Not enough data |
| | 0 to $1.00 \times 10^{-7}$ | $-\infty$ to -7 | $-\infty$ to 0.0000001 | 746 | 0 | 0 | Correct | Not enough data |
| **D** | Posterior probability range Pr<br>($H_p\vert LR$) | Log<sub>10</sub>LR<br>interval (2dp) | LR interval | $H_d$-true donors<br>Count | $H_p$-true donors<br>Count | Observed relative frequency of<br>$H_p$-true donors | Comment regarding $H_p$-true<br>donors | Interpretation |
| STRmix | 0.999999 to 1 | 6 to ∞ | 1001592 to ∞ | 0 | 922 | 1 | Correct | Not enough data |
|  | 0.99999 to 0.999999 | 5 to 6 | 100158 to 1001592 | 0 | 52 | 1 | More | Not enough data |
|  | 0.9999 to 0.99999 | 4 to 5 | 10015 to 100158 | 0 | 71 | 1 | More | Not enough data |
|  | 0.999 to 0.9999 | 3 to 4 | 1001 to 10015 | 0 | 54 | 1 | More | Not enough data |
|  | 0.99 to 0.999 | 2 to 3 | 99 to 1001 | 1 | 58 | 0.98 | Fewer | Anti-conservative |
|  | 0.95 to 0.99 | 1.28 to 2 | 19 to 99 | 0 | 30 | 1 | More | Conservative |

|  |  |  |  |  |  |  |  |
| --- | --- | --- | --- | --- | --- | --- | --- |
| 0.75 to 0.95 | 0.48 to 1.28 | 3 to 19 | 9 | 28 | 0.76 | Correct | Expected |
| 0.25 to 0.75 | -0.48 to 0.48 | 0.333 to 3 | 22 | 27 | 0.55 | Correct | N/A |
| 0.05 to 0.25 | -1.28 to -0.48 | 0.0527 to 0.333 | 63 | 6 | 0.087 | Correct | Expected |
| 0.01 to 0.05 | -2 to -1.28 | 0.0101 to 0.0527 | 61 | 5 | 0.076 | More | Anti-conservative |
| 0.001 to 0.01 | -3 to -2 | 0.001 to 0.0101 | 85 | 2 | 0.023 | More | Anti-conservative |
| $1.00 \times 10^{-4}$ to $1.00 \times 10^{-3}$ | -4 to -3 | 0.0001 to 0.001 | 70 | 0 | 0 | Fewer | Not enough data |
| $1.00 \times 10^{-5}$ to $1.00 \times 10^{-4}$ | -5 to -4 | 0.00001 to 0.0001 | 60 | 0 | 0 | Fewer | Not enough data |
| $1.00 \times 10^{-7}$ to $1.00 \times 10^{-5}$ | -7 to -5 | 0.0000001 to 0.00001 | 98 | 0 | 0 | Fewer | Not enough data |
| 0 to $1.00 \times 10^{-7}$ | $-\infty$ to -7 | $-\infty$ to 0.0000001 | 788 | 0 | 0 | Correct | Not enough data |

<sup>a</sup> Grey text = LR below $\log_{10} 0$ (approximate due to rounding).
<sup>b</sup> Not enough data = in theory, more $H_P$ -true counts in this range are needed to observe an $H_A$ -true result/ more $H_A$ -true counts in this range are needed to be able to observe an $H_P$ -true result.
<sup>c</sup> Anti-conservative = more false-positives were observed than expected based on the counts of $H_P$ -true donors in this range/ more false negatives were observed than expected based on the counts of $H_A$ -true donors in this range.
<sup>d</sup> Conservative (too close to LR=1) = with the number of $H_P$ -true counts in this range a false-positive would be expected but was not observed/ with the number of $H_A$ -true counts in this range a false-negative would be expected but was not observed.
<sup>e</sup> Expected = The observed relative frequency of $H_P$ -true donors falls within the posterior probability range.

Comparing calibration performance between the models, and for LR ranges >1, numbers of observed relative frequency values that fell within the posterior probability ranges were highest for HMC, then STRmix, followed by DNAStatistX and EuroForMix (Table 5). The lower performance of DNAStatistX and EuroForMix can be attributed to the H_d_-true scenarios yielding LRs that tend towards neutral evidence (Table 5AB, see bulge of H_d_-true counts in the log_10_LR interval –1.28 to +0.48).

Using the MLE model, the original dataset and this calibration metric for log_10_LRs >0, miscalibration is observed below ∼log_10_LR 3 when using θ = 0.01 (demonstrated through DNAStatistX and EuroForMix). Miscalibration in the range log_10_LR 2-3 was also observed for HMC and STRmix. The MLE model with θ = 0.03 shows miscalibration below ∼log_10_LR 2 (demonstrated through DNAStatistX).

Importantly, the poorer results observed in Table 5A compared to Table 4A can be attributed to the smaller dataset size. DNAStatistX θ = 0.01 and θ = 0.03 demonstrated better calibration performance with the extended dataset in Table 4A. Interestingly, we also observe slight variations in calibration performance across different STRmix datasets in the current study (Table 5D), Bright et al. [29] and Buckleton et al. [18]. This may be attributed to differences in dataset size, ratio of H_d_-true to H_p_-true scenarios and/or posterior probability ranges.

Furthermore, although the calibration tables provide an insight into calibration performance, it must be noted that these metrics have a few drawbacks, apart from the sensitivity to database size. For example, the relative frequency of H_p_-true donors in posterior probability ranges were, in some cases, on the edge of the posterior probability range and with a slight change (or just rounding) an anti-conservative result then fell into the category ‘expected’ or ‘conservative’. This was observed for almost all of the software. Furthermore, the extent to which an LR is over-or understated cannot be read from the calibration tables, unlike in other calibration plot metrics (discussed below).

### 3.3 Calibration plots

Figs. 8 and 9 show calibration results presented in 1) ECE plots with Cllr and Cllr*^cal^* values, 2) PAV plots with devPAV values and 3) fiducial calibration discrepancy plots. Fig. 8 shows DNAStatistX results when using the extended dataset, Fig. 9 shows DNAStatistX and EuroForMix results when using the original dataset.

**Figure 8.**
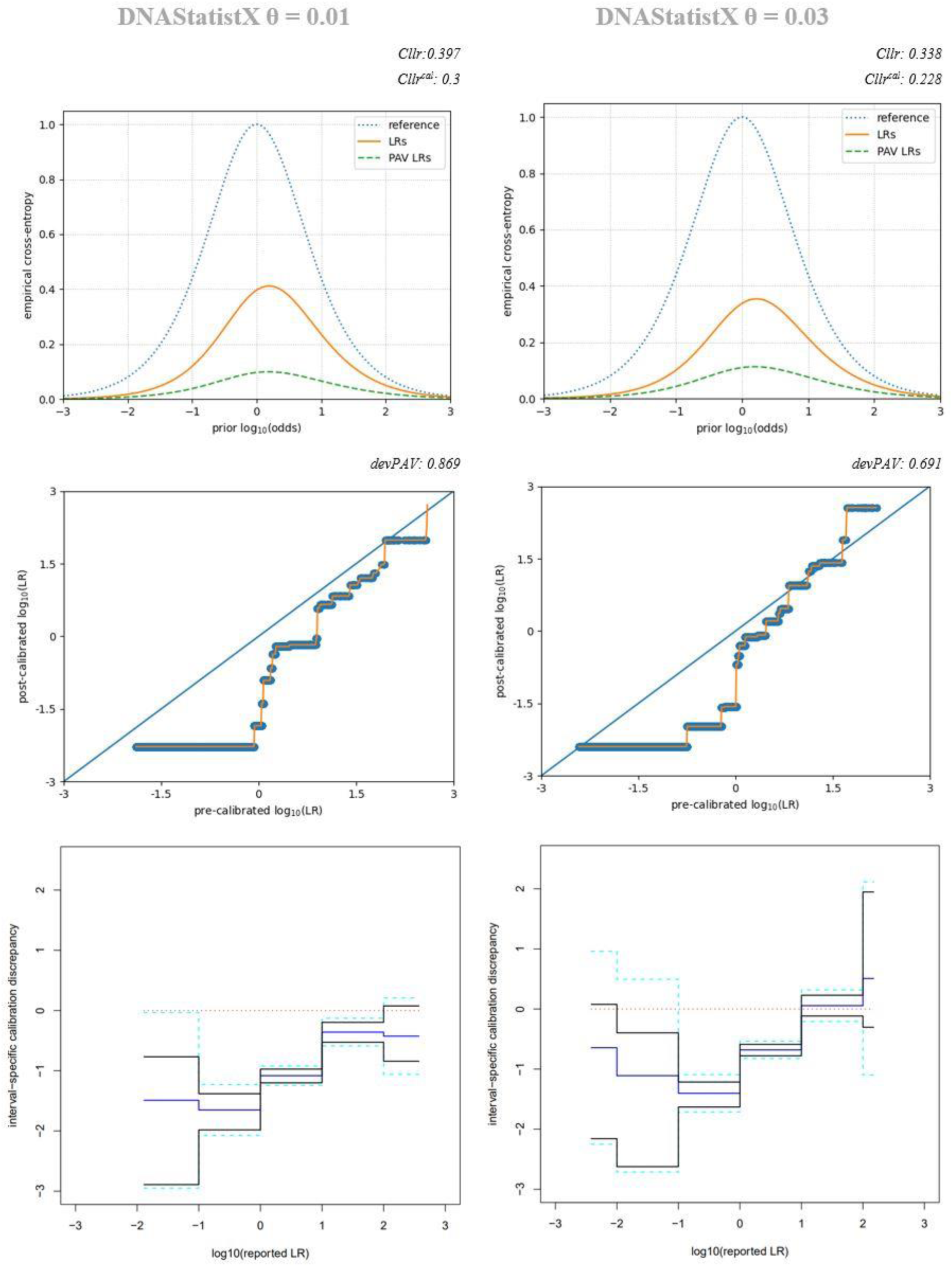
Calibration plots of DNAStatistX using the extended dataset. Top: ECE plots and Cllr and Cllr*^cal^* values (3dp). Middle: PAV plots and devPAV values (3dp). Bottom: Fiducial calibration discrepancy plots.

**Figure 9.**
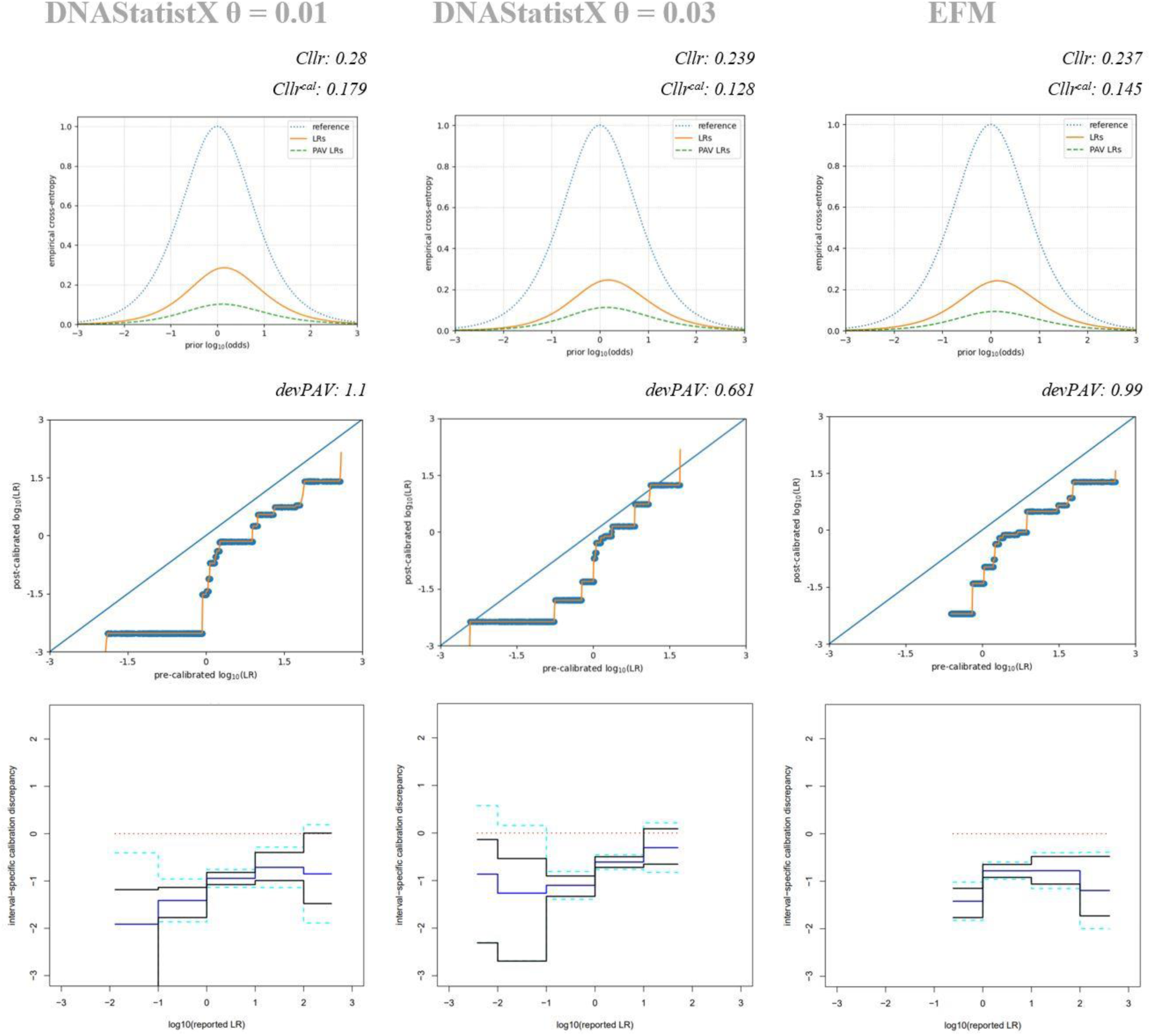
Calibration plots of DNAStatistX and EuroForMix using the original dataset. Top: ECE plots and Cllr and Cllr*^cal^* values (3dp). Middle: PAV plots and devPAV values (3dp). Bottom: Fiducial calibration discrepancy plots.

The ECE plots demonstrate that the value of ECE is lower than the value of the neutral method, and thus shows that the method provides useful information for evidence evaluation. This holds true for all values of the prior-log-odds, for all PG software.

With respect to the extended DNAStatistX dataset, the ECE plots in Fig. 8 show lower values for Cllr and Cllr*^cal^* when using DNAStatistX with θ =0.03 than θ =0.01. This confirms the findings discussed in the previous sections, i.e. that DNAStatistX shows better calibration performance with θ =0.03 than with θ =0.01. This observation is further substantiated with the fiducial calibration discrepancy plots (data more close to horizontal line at value 0 when using θ =0.03) and the PAV plots (datapoints more close to diagonal line when using θ =0.03). In particular, with the extended DNAStatistX θ =0.03 dataset, at log10LRs above 1, the calibration discrepancy followed the perfect calibration lines closely and calibration performance was comparable to HMC and STRmix (Supplementary Fig. 3). Notably, LRs were tending towards understatement between log_10_LRs +2 and +3.

Then, with the original dataset, the PAV and fiducial calibration plots again indicate that low LRs for DNAStatistX and EuroForMix (both θ = 0.01) are overstated with varying degrees, depending on the LR range. When LRs were calculated in DNAStatistX with θ = 0.03 and using the original dataset, this trend of LR overstatement was observed but to a lesser degree (Fig. 9). Consistent with observations in the calibration tables, better calibration performance was observed using the extended DNAStatistX dataset (Fig. 8) than the original dataset (Fig. 9) in the calibration discrepancy plots and PAV plots. Interestingly, according to the ECE plots, the opposite was observed: calibration appeared better using the DNAStatistX original dataset. This may be explained by the Cllr taking account of all LRs, including the very small and very large ones. On the contrary, the devPAV only considers LRs in the range where H_p_-true and H_d_-true LRs overlap. The extended dataset includes additional H_d_-true LRs, and these were computed only for scenarios that yielded low H_p_-true LRs. Due to these challenging samples, many non-contributor LRs tended to fall around 1. Consequently, adding H_d_-true LRs that fall around 1 may improve the devPAV, while they make the Cllr worse as the proportion of H_d_-true LRs below the smallest H_p_-true LR decreases. We note however, that much more detailed examination of calibration results from the original dataset can be performed using the fiducial calibration discrepancy plots and the PAV plots than ECE. I.e. fiducial calibration discrepancy plots and PAV plots consider ranges in which H_p_-true and H_d_-true LRs overlap and provide information on the extent to which LRs in these ranges are over-or understated.

The original, and smaller, dataset further enables comparison between models. Fig. 9 shows that the calibration performance of DNAStatistX and EuroForMix are quite comparable for the θ =0.01 results, with respect to trends of LR overstatement. However, some differences in LRs between the software can be expected, since a slightly different approach was used for model selection in this study (see section 2.2.1). Supplementary Fig. 3 shows HMC and STRmix plots, which demonstrate better calibration performance in the lower LR ranges than the models incorporating the MLE method (Figs. 8-9, θ = 0.01 data), through lower Cllr, Cllr*^cal^* and devPAV values.

Altogether, we have observed evidence of miscalibration of low LRs in the calibration plots, when using the MLE method in DNAStatistX/EuroForMix. Still, calibration performance improved when using θ = 0.03, above log_10_LR ∼1, and when the extended dataset was used, the discrepancies were further reduced (Fig. 8-9).

### 3.6 Turing expectation

Using Turing expectation tests (non-contributor tests), LRs of true– and non-contributor scenarios were compared. For a given true-contributor scenario, the LR_*x*_ was compared to 500 LRs computed under the same model parameters as the true-contributor scenario, but with random non-contributors set as the POI. In theory, the fraction of non-contributor scenarios producing an LR ≥ LR_*x*_ is expected to be *at most* 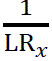. If this is true, i.e. the distribution is within predicted limits given Turing’s bound, the Turing expectation test was scored passed. Supplementary Table 3 provides a summary of the DNAStatistX H_p_-true scenarios for which Turing tests were performed, including mixture profile and POI characteristics. Fig. 10 shows that all of the H_p_-true scenario results (ordered from highest to lowest H_p_-true scenario LR) passed the Turing expectation (both with θ = 0.01 and θ = 0.03). This is represented by black horizontal lines indicating the percentage of the expected numbers of H_d_-true LRs ≥ LR_*x*_ that were actually observed. For all scenarios, the number of observed H_d_-true LRs ≥ LR_*x*_ never exceeded the expected numbers, even if there was overlap of H_p_-true and H_d_-true LRs. However, a limitation must be noted that when the log_10_LR was <0, the expected number of violations exceeded the N of 500 (Supplementary Figs. 4-5). Although none of the observed violations reached 500, some were very close (scenario 21 in Fig. 10A and scenario 26 in Fig. 10B).

**Figure 10.**
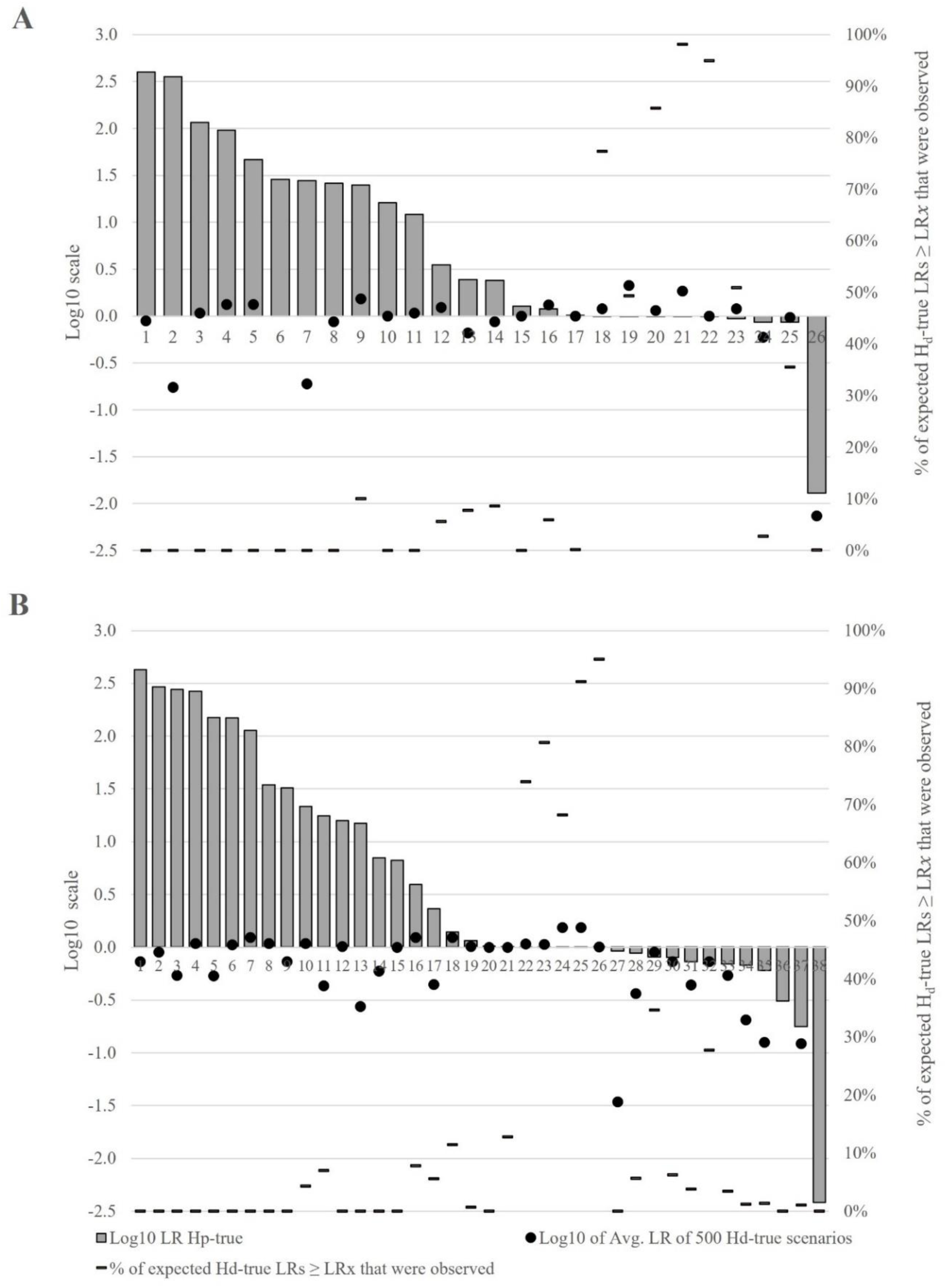
Turing non-contributor test results. A) DNAStatistX θ = 0.01, B) DNAStatistX θ = 0.03. Data points ordered from highest to lowest H_p_-true scenario log_10_LR (LR*x*) (grey bars, left y-axes). The black dots represent the log_10_ of the average LR of the 500 H_d_-true scenarios to which each respective H_p_-true scenario LR*x* is being compared (left y-axes). The black horizontal lines represent the percentage of the expected numbers of Hd-true LRs ≥ LR*x* that were actually observed (right y-axes).

As an additional layer to testing, we analysed whether Turing expectation would pass for the non-contributor scenarios. This meant, one by one, each non-contributor POI scenario was treated as the ‘reference LR’ scenario and the LRs from the other 499 non-contributors for the same profile scenario were used for comparison. In line with the above limitation, we analysed this until the number of LRs ≥ LR*x* expected under Turing expectation exceeded 499. Interestingly, all of the non-contributor scenarios that we analysed passed Turing expectation (Supplementary Table 4). Furthermore, the numbers of expected LRs ≥ LR*x* under Turing expectation increased at a much higher rate than the observed numbers of LRs ≥ LR*x*. Therefore, one could speculate that the Turing expectation would continue to pass beyond this point.

Overall, the numbers of H_d_-true test LRs that exceeded the H_p_-true LRs were clearly lower than expected from Turing’s rule, regardless of the considered Fst value. The DNAStatistX results confirm previous findings using other PG software demonstrating the expectation that a portion of non-contributors will have an LR >1 (Supplementary Fig. 5-6) but that the distribution is well within predicted limits given Turing’s bound [28]. Turing expectation tests, Tippett tests, or non-contributor performance tests have been suggested to perform on a per case basis to gain insight in the robustness of the LR measurement itself [34]. Following Turing, the DNAStatistX LRs obtained in this study are regarded robust and in [28] such results were previously taken as strong evidence that low LRs are reliable.

Another measure for robustness is examining whether the log_10_ of the average of the H_d_-true LRs is close to one, although authors of [12] reiterate that this is more an indication of the performance of the models used to generate the LR than of the LR itself. This was true for many, but not for all, DNAStatistX results in this study (Fig. 10, most, but not all are close to 0), and thus low LRs in DNAStatistX do not appear as robust under this metric.

## 4. Discussion and conclusions

Previous studies have reported on PG systems validity according to guidelines for developmental and internal validation of PG software. These guidelines mainly focus on discriminatory power. Except for Tippett plots and calibration tables, very few studies have reported on calibration metrics for PG system validation [26, 27, 29]. The current study assessed the discriminatory power and calibration performance of PG software in LR ranges <10,000, with focus on DNAStatistX and EuroForMix results. Therefore, the performance of the MLE method was evaluated in low LR ranges.

From the results of the current study, we aimed to interrogate and make recommendations with respect to lower range LRs obtained using DNAStatistX and EuroForMix and minimum reporting thresholds. Many (European) laboratories apply a lower threshold for reporting and therefore overstated low LRs have no effect for reporting. LRs below the threshold are not reported and can be considered ‘neutral’ evidence. Setting the LR threshold arbitrarily high, however, wastes relevant evidential value. Therefore, the use of a lower threshold for reporting is argued and discouraged [10, 11]. If a PG software is perfectly calibrated throughout the entire LR range, a lower LR threshold should not be needed. The reason for applying an LR threshold and the height of this threshold can be multifaceted, though (as to our knowledge) no literature exists that substantiates a laboratory’s LR threshold by calibration results as presented in this study.

In the current study, as expected, DNAStatistX and EuroForMix exhibited the most similar performance amongst the PG software tested. Furthermore, for these PG software, good discriminatory power was previously presented [8, 14, 15, 17, 20, 21, 23] and confirmed in this study (e.g. Table 3). The discriminatory performance was perfect above log_10_LR +3 using θ =0.01, which was also observed for HMC and STRmix (Table 3). Using DNAStatistX with θ = 0.03, perfect discriminatory power was observed above log_10_LR +2.

Regarding calibration, the calibration tables and various calibration plots demonstrate a dependency of Fst value and dataset size on calibration performance (Tables 4-5; Fig. 8-9). Miscalibration of LRs >1 (log_10_LRs >0) was observed using the MLE-based models, specifically with the original dataset and lower value for Fst (θ = 0.01): LRs up to ∼1,000 (log_10_ LR 3) tend to be overstated. Yet when using the extended dataset and θ = 0.03, calibration performance appeared to be satisfactory above log_10_LR 0, and very similar to PG systems HMC and STRmix. If anything, there was slight LR understatement approaching as LRs increased, which is more favorable than overstatement, i.e. closer to neutral evidence (Table 4; Fig. 8; Supplementary Fig. 3). The results confirm previous observations [15, 16, 17] that H_d_-true scenario LRs can be much larger (i.e. tend towards neutral evidence) in DNAStatistX and EuroForMix than LRs obtained using other PG software (Tables 4-5; Fig. 8-9). This overestimation may be attributed to the nature of the MLE model, where the likelihood is maximised under H_d_ as well as H_p_ [8, 39]. Previously, this has also been reported as a cause of differences in ‘low’ LRs between EuroForMix and STRmix [18]. Most importantly, the results confirm previous observations [15, 19, 20, 21] that high H_p_-true LRs (LR >1,000) were similar to other PG software when performed with the same input data and propositions (Fig. 4).

We employed a variety of metrics to assess performance, both well-known and relatively novel for PG evaluation. In our experience, the PAV and fiducial calibration discrepancy plots showed the most detailed information on calibration performance; with the confidence intervals provided by the fiducial calibration discrepancy plots providing an additional layer of insight. It should be emphasised again that calibration can only be evaluated when a dataset is sufficiently large with respect to the LR range. Last and surprisingly, DNAStatistX Turing expectation results always passed, indicating that low LRs were well-calibrated and robust (Table 6; Supplementary Table 4). In light of the other results, this calls into question the utility of Turing expectation tests when deciding whether a low LR is robust, and how these tests indicate calibration. They appear a poor test of calibration. We conclude that Turing expectation tests are most useful to explain to courts generally how the LR performs with respect to mathematical expectations of probabilistic models.

As mentioned previously, if a model is well calibrated throughout the entire LR ranges, a lower LR threshold for reporting is redundant. Furthermore, the use of minimum reporting thresholds are increasingly discouraged. However, due to evidence of miscalibration in low LRs ranges in the current study, we would recommend caution if/when reporting LRs below log_10_LR +3 (LR 1,000) if an MLE model is used with θ = 0.01. Indeed, there is risk of LR overestimation. However, raising the Fst to theta=0.03 appears to decrease the risk when log_10_LRs > 0 (i.e. LR > 1). This simple expedient produces calibration results that are comparable to those achieved by HMC/ STRmix for this range. Of course, a laboratory’s choice for Fst values is dependent on the relevant population database for a case as well as the results of internal validation studies, though the effect of the value for Fst on calibration performance is relevant to present to end-users as an option to explore. For labs wishing to report LR<1,000, we suggest that application of Fst=0.03 is an option to consider.

Regardless of the software, above their respective recommended minimum reporting thresholds (or even if none is selected), all LRs must be well-calibrated and robust as per mathematical expectation. Furthermore, model validation provides an additional safeguard for DNAStatistX/ EuroForMix, as this must also pass for the LR to be reported.

The international relevance of this research should be mentioned. Most EU countries do not report H_d_-supporting LR values. This is because it would be unusual for low LRs that do not provide strong support for a prosecution proposition to be presented in court. If one wishes to use no LR threshold or set it as low as possible (without the risk of reporting over-or understated LRs), we emphasise the need to analyse both the discriminatory power and calibration performance of PG software to justify decisions regarding LR reporting and we reiterate the suggestion that Fst=0.03 when if it is desired to report LR<1,000.

As a final note we would like to emphasise that it is not just an LR threshold that defines whether a weight of evidence is reported. Laboratories may limit calculations based on e.g. the estimated NoC, the number of unknown contributors, or the number of minor contributors [40]. In addition, differences in choices for e.g. STR typing kit, analytical thresholds and Fst value yield differences in LRs and may result in an LR that will or will not exceed the LR threshold, or not be computed at all. Efforts to decrease laboratory differences are encouraged, though it is not realistic that these will be eliminated completely. This emphasises the importance of transparency to help form a basis to make informed choices for laboratory practice. This study provides an example.

## Acknowledgements

We thank Jerry Hoogenboom and Klaas Slooten for useful discussions on the project. We are grateful to Jerry, for important help with the data, data format and command-line. Klaas and Jan Hannig are thanked for providing insight in the fiducial calibration discrepancy plots. Lastly, we extend thanks to Mateusz Susik and, Ivo F. Sbalzarini for sharing their dataset.

## Notes

### Competing Interest Statement

Authors are developers and/or employed at the institutes developing EuroForMix or DNAStatistX, but there is no financial interest in the software.

