## Supplementary Tables 2-4 for "‘Low’ LRs obtained from DNA mixtures: On calibration and discrimination performance of probabilistic genotyping software"

Supplementary Table 2. Sensitivity, specificity, PPV, NPV and accuracy results for different PG software (3dp). Results for two- (A), three- (B) or four-person mixtures (C). Color gradients set within each variable; highest rates are shown on a scale from dark green, to light green, to yellow, to orange, to red.

| **A** | Software | | Sensitivity | Specificity | Accuracy | PPV | NPV |
|---|---|---|---|---|---|---|---|
| | HMC [17] | | 0.997 | 1 | 0.998 | 1 | 0.997 |
| | STRmix [17] | | 0.987 | 0.997 | 0.992 | 0.997 | 0.987 |
| | EFM | | 0.994 | 0.984 | 0.989 | 0.984 | 0.994 |
| | DNAStatistX | θ = 0.01 | 0.994 | 0.957 | 0.975 | 0.957 | 0.994 |
| | | θ = 0.03 | 0.99 | 0.957 | 0.973 | 0.957 | 0.99 |
| **B** | Software | | Sensitivity | Specificity | Accuracy | PPV | NPV |
| | HMC [17] | | 0.978 | 0.995 | 0.987 | 0.995 | 0.978 |
| | STRmix [17] | | 0.974 | 0.995 | 0.984 | 0.995 | 0.974 |
| | EFM | | 0.991 | 0.919 | 0.954 | 0.919 | 0.991 |
| | DNAStatistX | θ = 0.01 | 0.991 | 0.858 | 0.92 | 0.858 | 0.991 |
| | | θ = 0.03 | 0.987 | 0.88 | 0.93 | 0.88 | 0.987 |
| **C** | Software | | Sensitivity | Specificity | Accuracy | PPV | NPV |
| | HMC [17] | | 0.981 | 0.979 | 0.98 | 0.979 | 0.981 |
| | STRmix [17] | | 0.981 | 0.973 | 0.977 | 0.973 | 0.981 |
| | EFM | | 0.994 | 0.795 | 0.883 | 0.795 | 0.994 |
| | DNAStatistX | θ = 0.01 | 0.994 | 0.703 | 0.823 | 0.703 | 0.994 |
| | DNAStatistX | θ = 0.03 | 0.984 | 0.757 | 0.856 | 0.757 | 0.984 |

### Supplementary Table 3. Details of the $H_p$-true scenario details for which Turing expectation tests were performed.

| Profile | Log10LR | LR | Selected model condition | Total Amount (pg) | Ratio | Treatment | POI | Type of POI |
|---|---|---|---|---|---|---|---|---|
| F10_RD14-0003-39_40-1_2-M3c-0.045GF-Q1.0_06.15sec | -0.0685 | 0.85416 | SM OFF | 45 | 1;2 | c | 39 | Minor |
| G11_RD14-0003-39_40-1_2-M4e-0.045GF-Q2.4_07.15sec | -0.0065 | 0.9851 | SM OFF | 45 | 1;2 | e | 39 | Minor |
| A09_RD14-0003-49_50_29-1_4_1-M3e-0.186GF-Q1.6_01.15sec | -0.0284 | 0.9368 | SM OFF | 186 | 1;4;1 | e | 29 | Minor |
| B07_RD14-0003-49_50_29-1_4_1-M3U105-0.09GF-Q6.1_02.15sec | -0.0003 | 0.99924 | BW | 90 | 1;4;1 | U105 | 29 | Minor |
| D08_RD14-0003-30_31_32-1_4_4-M2U105-0.135GF-Q5.1_04.15sec | -0.0679 | 0.85523 | SM OFF | 135 | 1;4;4 | U105 | 30 | Minor |
| F03_RD14-0003-49_50_29-1_4_1-M3S30-0.09GF-Q2.9_06.15sec | -0.0011 | 0.99742 | SM OFF | 90 | 1;4;1 | S30 | 29 | Minor |
| E05_RD14-0003-33_34_35_36-1_4_4_1-M2a-0.63GF-Q0.5_05.15sec | -1.8894 | 0.0129 | FW | 630 | 1;4;4;1 | a | 33 | Minor |
| F05_RD14-0003-44_45_46_47-1_1_4_1-M2S30-0.217GF-Q3.3_06.15sec | -0.0002 | 0.99952 | SM OFF | 217 | 1;1;4;1 | S30 | 44 | Minor |
| G05_RD14-0003-44_45_46_47-1_1_4_1-M2S30-0.105GF-Q3.3_07.15sec | -0.0002 | 0.99943 | FW | 105 | 1;1;4;1 | S30 | 44 | Minor |
| B04_RD14-0003-40_41-1_4-M4e-0.075GF-Q0.7_02.15sec | 2.6016 | 399.576 | BW & FW | 75 | 1;4 | e | 40 | Minor |
| D01_RD14-0003-44_45-1_1-M4I15-0.03GF-Q1.2_04.15sec | 1.20822 | 16.1518 | FW | 30 | 1;1 | I15 | 45 | Equal |
| D03_RD14-0003-40_41-1_4-M3U105-0.075GF-Q14.8_04.15sec | 2.55282 | 357.122 | SM OFF | 75 | 1;4 | U105 | 40 | Minor |
| G02_RD14-0003-31_32-1_1-M1e-0.062GF-Q14.4_07.15sec | 1.44125 | 27.6218 | SM OFF | 62 | 1;1 | e | 31 | Equal |
| A03_RD14-0003-36_37_38-1_2_1-M3e-0.06GF-Q1.6_01.15sec | 1.08308 | 12.1081 | SM OFF | 60 | 1;2;1 | e | 36 | Minor |
| A03_RD14-0003-36_37_38-1_2_1-M3e-0.06GF-Q1.6_01.15sec | 2.06274 | 115.542 | SM OFF | 60 | 1;2;1 | e | 38 | Minor |
| B05_RD14-0003-41_42_43-1_9_1-M2e-0.165GF-Q1.7_02.15sec | 0.07563 | 1.19023 | BW | 165 | 1;9;1 | e | 43 | Minor |
| F05_RD14-0003-46_47_48-1_1_1-M3U60-0.045GF-Q4.0_06.15sec | 0.38772 | 2.44184 | SM OFF | 45 | 1;1;1 | U60 | 48 | Equal |
| G07_RD14-0003-30_31_32-1_4_4-M4I22-0.135GF-Q4.7_07.15sec | 0.5459 | 3.5148 | BW | 135 | 1;4;4 | I22 | 30 | Minor |
| H06_RD14-0003-49_50_29-1_4_1-M4I35-0.09GF-QLAND_08.15sec | 0.1053 | 1.27439 | FW | 90 | 1;4;1 | I35 | 29 | Minor |
| H06_RD14-0003-49_50_29-1_4_1-M4I35-0.09GF-QLAND_08.15sec | 0.00631 | 1.01462 | FW | 90 | 1;4;1 | I35 | 49 | Minor |
| C07_RD14-0003-33_34_35_36-1_4_4_1-M2e-0.15GF-Q0.9_03.15sec | 0.38016 | 2.39974 | SM OFF | 150 | 1;4;4;1 | e | 33 | Minor |
| C07_RD14-0003-33_34_35_36-1_4_4_1-M2e-0.15GF-Q0.9_03.15sec | 1.41581 | 26.0503 | SM OFF | 150 | 1;4;4;1 | e | 36 | Minor |
| E05_RD14-0003-36_37_38_39-1_2_2_1-M2e-0.378GF-Q1.2_05.15sec | 1.66657 | 46.4059 | BW | 378 | 1;2;2;1 | e | 39 | Minor |
| G07_RD14-0003-37_38_39_40-1_9_9_1-M2a-0.75GF-Q0.4_07.15sec | 1.45683 | 28.6306 | FW | 750 | 1;9;9;1 | a | 37 | Minor |
| H01_RD14-0003-40_41_42_43-1_1_1_1-M4d-0.124GF-Q1.7_08.15sec | 1.98058 | 95.626 | SM OFF | 124 | 1;1;1;1 | d | 42 | Equal |
| H03_RD14-0003-44_45_46_47-1_1_4_1-M2e-0.441GF-Q1.5_08.15sec | 1.39864 | 25.0404 | BW & FW | 441 | 1;1;4;1 | e | 44 | Minor |
| B08_RD14-0003-37_38_39_40-1_9_9_1-M2a-0.3GF-Q0.4_02.15sec | 3.42212 | 2643.11 | BW | 300 | 1;9;9;1 | a | 37 | Minor |
| A03_RD14-0003-40_41-1_4-M4a-0.075GF-Q0.6_01.15sec | -0.51137 | 0.308054 | SM OFF | 75 | 1;4 | a | 40 | Minor |
| F10_RD14-0003-39_40-1_2-M3c-0.045GF-Q1.0_06.15sec | -0.15851 | 0.694201 | SM OFF | 45 | 1;2 | c | 39 | Minor |
| G11_RD14-0003-39_40-1_2-M4e-0.045GF-Q2.4_07.15sec | -0.00425 | 0.990251 | SM OFF | 45 | 1;2 | e | 39 | Minor |
| A08_RD14-0003-30_31_32-1_4_4-M2U60-0.135GF-Q2.7_01.15sec | -0.2181 | 0.605204 | SM OFF | 135 | 1;4;4 | U60 | 30 | Minor |
| A09_RD14-0003-49_50_29-1_4_1-M3e-0.186GF-Q1.6_01.15sec | -0.09195 | 0.809187 | SM OFF | 186 | 1;4;1 | e | 29 | Minor |
| B07_RD14-0003-49_50_29-1_4_1-M3U105-0.09GF-Q6.1_02.15sec | -0.00028 | 0.999363 | BW | 90 | 1;4;1 | U105 | 29 | Minor |
| B09_RD14-0003-49_50_29-1_4_1-M3e-0.09GF-Q1.6_02.15sec | -0.05361 | 0.883881 | SM OFF | 90 | 1;4;1 | e | 49 | Minor |
| D08_RD14-0003-30_31_32-1_4_4-M2U105-0.135GF-Q5.1_04.15sec | -0.75128 | 0.177305 | SM OFF | 135 | 1;4;4 | U105 | 30 | Minor |
| F03_RD14-0003-49_50_29-1_4_1-M3S30-0.09GF-Q2.9_06.15sec | -0.0011 | 0.997465 | SM OFF | 90 | 1;4;1 | S30 | 29 | Minor |
| A03_RD14-0003-50_29_30_31-1_1_2_1-M3e-0.075GF-Q1.1_01.15sec | -0.09547 | 0.802652 | SM OFF | 75 | 1;1;2;1 | e | 31 | Minor |
| C11_RD14-0003-48_49_50_29-1_4_4_4-M3d-0.195GF-Q0.9_03.15sec | -0.16836 | 0.678646 | SM OFF | 195 | 1;4;4;4 | d | 48 | Minor |
| E01_RD14-0003-50_29_30_31-1_1_2_1-M3a-0.075GF-Q0.5_05.15sec | -0.03415 | 0.924368 | SM OFF | 75 | 1;1;2;1 | a | 31 | Minor |
| E04_RD14-0003-48_49_50_29-1_4_4_4-M2U105-0.195GF-Q12.1_05.15sec | -0.13618 | 0.730842 | BW | 195 | 1;4;4;4 | U105 | 48 | Minor |
| E05_RD14-0003-33_34_35_36-1_4_4_1-M2a-0.63GF-Q0.5_05.15sec | -2.41763 | 0.003823 | FW | 630 | 1;4;4;1 | a | 33 | Minor |
| F05_RD14-0003-44_45_46_47-1_1_4_1-M2S30-0.217GF-Q3.3_06.15sec | -0.00025 | 0.999424 | SM OFF | 217 | 1;1;4;1 | S30 | 44 | Minor |
| G05_RD14-0003-44_45_46_47-1_1_4_1-M2S30-0.105GF-Q3.3_07.15sec | -0.00102 | 0.997655 | FW | 105 | 1;1;4;1 | S30 | 44 | Minor |
| G06_RD14-0003-33_34_35_36-1_4_4_1-M2d-0.15GF-Q0.9_07.15sec | -0.15919 | 0.69312 | FW | 150 | 1;4;4;1 | d | 33 | Minor |
| A02_RD14-0003-40_41-1_4-M3S30-0.075GF-Q4.0_01.15sec | 1.174662 | 14.95072 | SM OFF | 75 | 1;4 | S30 | 40 | Minor |
| B04_RD14-0003-40_41-1_4-M4e-0.075GF-Q0.7_02.15sec | 2.177234 | 150.3951 | BW & FW | 75 | 1;4 | e | 40 | Minor |
| D01_RD14-0003-44_45-1_1-M4I15-0.03GF-Q1.2_04.15sec | 0.82118 | 6.62491 | FW | 30 | 1;1 | I15 | 45 | Equal |
| E02_RD14-0003-31_32-1_1-M2d-0.03GF-Q6.0_05.15sec | 1.201058 | 15.8876 | SM OFF | 30 | 1;1 | d | 31 | Equal |
| E02_RD14-0003-31_32-1_1-M2d-0.03GF-Q6.0_05.15sec | 0.063413 | 1.157213 | SM OFF | 30 | 1;1 | d | 32 | Equal |
| A02_RD14-0003-36_37_38-1_2_1-M4c-0.06GF-Q1.1_01.15sec | 1.508772 | 32.268 | SM OFF | 60 | 1;2;1 | c | 36 | Minor |
| A02_RD14-0003-36_37_38-1_2_1-M4c-0.06GF-Q1.1_01.15sec | 2.629497 | 426.0861 | SM OFF | 60 | 1;2;1 | c | 38 | Minor |
| D05_RD14-0003-46_47_48-1_1_1-M4I22-0.045GF-Q1.4_04.15sec | 2.426765 | 267.1563 | SM OFF | 45 | 1;1;1 | I22 | 46 | Equal |
| D05_RD14-0003-46_47_48-1_1_1-M4I22-0.045GF-Q1.4_04.15sec | 1.54048 | 34.71204 | SM OFF | 45 | 1;1;1 | I22 | 47 | Equal |
| D05_RD14-0003-46_47_48-1_1_1-M4I22-0.045GF-Q1.4_04.15sec | 1.332267 | 21.49151 | SM OFF | 45 | 1;1;1 | I22 | 48 | Equal |
| F04_RD14-0003-41_42_43-1_9_1-M2d-0.165GF-Q1.2_06.15sec | 2.171888 | 148.5554 | BW & FW | 165 | 1;9;1 | d | 43 | Minor |
| F09_RD14-0003-47_48_49-1_9_9-M2e-0.5GF-Q1.5_06.15sec | 1.245382 | 17.5947 | BW | 500 | 1;9;9 | e | 47 | Minor |
| H06_RD14-0003-49_50_29-1_4_1-M4I35-0.09GF-QLAND_08.15sec | 0.010785 | 1.025145 | FW | 90 | 1;4;1 | I35 | 29 | Minor |
| H06_RD14-0003-49_50_29-1_4_1-M4I35-0.09GF-QLAND_08.15sec | 0.000189 | 1.000436 | FW | 90 | 1;4;1 | I35 | 49 | Minor |
| D01_RD14-0003-40_41_42_43-1_1_1_1-M4I15-0.06GF-Q1.6_04.15sec | 2.466379 | 292.6702 | SM OFF | 60 | 1;1;1;1 | I15 | 41 | Equal |
| D09_RD14-0003-37_38_39_40-1_9_9_1-M2e-0.62GF-Q0.8_04.15sec | 0.366809 | 2.32707 | BW & FW | 620 | 1;9;9;1 | e | 37 | Minor |
| E05_RD14-0003-36_37_38_39-1_2_2_1-M2e-0.378GF-Q1.2_05.15sec | 0.84659 | 7.024087 | BW | 378 | 1;2;2;1 | e | 39 | Minor |
| G06_RD14-0003-33_34_35_36-1_4_4_1-M2d-0.15GF-Q0.9_07.15sec | 2.442774 | 277.1879 | FW | 150 | 1;4;4;1 | d | 36 | Minor |
| H02_RD14-0003-44_45_46_47-1_1_4_1-M3U60-0.105GF-Q28.7_08.15sec | 0.592668 | 3.914424 | BW | 105 | 1;1;4;1 | U60 | 44 | Minor |
| H02_RD14-0003-44_45_46_47-1_1_4_1-M3U60-0.105GF-Q28.7_08.15sec | 2.054762 | 113.4388 | BW | 105 | 1;1;4;1 | U60 | 45 | Minor |
| H02_RD14-0003-44_45_46_47-1_1_4_1-M3U60-0.105GF-Q28.7_08.15sec | 0.144309 | 1.394148 | BW | 105 | 1;1;4;1 | U60 | 47 | Minor |
| G02_RD14-0003-50_29_30_31-1_1_2_1-M2e-0.315GF-Q1.0_07.15sec | 3.36406 | 2312.4 | BW | 315 | 1;1;2;1 | e | 29 | Minor |
| G07_RD14-0003-37_38_39_40-1_9_9_1-M2a-0.75GF-Q0.4_07.15sec | 0.49605 | 3.13367 | FW | 750 | 1;9;9;1 | a | 37 | Minor |

### Supplementary Table 4. Turing $H_d$-true scenarios where Turing expectation was analysed and the results.

| Condition | NoC | Mixture profile | Number of non-contributor scenarios where N LRs ≥ LR*x* expected under Turing expectation was <499 | Selected model condition | % that passed Turing expectation |
|---|---|---|---|---|---|
| θ = 0.01 | 2 | B04_RD14-0003-40_41-1_4-M4e-0.075GF-Q0.7_02.15sec | 59 | BW & FW | 100% |
| | | D01_RD14-0003-44_45-1_1-M4I15-0.03GF-Q1.2_04.15sec | 237 | FW | 100% |
| | | D03_RD14-0003-40_41-1_4-M3U105-0.075GF-Q14.8_04.15sec | 15 | SM OFF | 100% |
| | | F10_RD14-0003-39_40-1_2-M3c-0.045GF-Q1.0_06.15sec | 175 | SM OFF | 100% |
| | | G02_RD14-0003-31_32-1_1-M1e-0.062GF-Q14.4_07.15sec | 21 | SM OFF | 100% |
| | | G11_RD14-0003-39_40-1_2-M4e-0.045GF-Q2.4_07.15sec | 293 | SM OFF | 100% |
| | 3 | A03_RD14-0003-36_37_38-1_2_1-M3e-0.06GF-Q1.6_01.15sec | 320 | SM OFF | 100% |
| | | A09_RD14-0003-49_50_29-1_4_1-M3e-0.186GF-Q1.6_01.15sec | 98 | SM OFF | 100% |
| | | B05_RD14-0003-41_42_43-1_9_1-M2e-0.165GF-Q1.7_02.15sec | 402 | BW | 100% |
| | | B07_RD14-0003-49_50_29-1_4_1-M3U105-0.09GF-Q6.1_02.15sec | 309 | BW | 100% |
| | | D08_RD14-0003-30_31_32-1_4_4-M2U105-0.135GF-Q5.1_04.15sec | 11 | SM OFF | 100% |
| | | F03_RD14-0003-49_50_29-1_4_1-M3S30-0.09GF-Q2.9_06.15sec | 397 | SM OFF | 100% |
| | | F05_RD14-0003-46_47_48-1_1_1-M3U60-0.045GF-Q4.0_06.15sec | 48 | SM OFF | 100% |
| | | G07_RD14-0003-30_31_32-1_4_4-M4I22-0.135GF-Q4.7_07.15sec | 288 | BW | 100% |
| | | H06_RD14-0003-49_50_29-1_4_1-M4I35-0.09GF-QLAND_08.15sec | 285 | FW | 100% |
| | 4 | C07_RD14-0003-33_34_35_36-1_4_4_1-M2e-0.15GF-Q0.9_03.15sec | 77 | SM OFF | 100% |
| | | E05_RD14-0003-33_34_35_36-1_4_4_1-M2a-0.63GF-Q0.5_05.15sec | 1 | FW | 100% |
| | | E05_RD14-0003-36_37_38_39-1_2_2_1-M2e-0.378GF-Q1.2_05.15sec | 353 | BW | 100% |
| | | F05_RD14-0003-44_45_46_47-1_1_4_1-M2S30-0.217GF-Q3.3_06.15sec | 291 | SM OFF | 100% |
| | | G05_RD14-0003-44_45_46_47-1_1_4_1-M2S30-0.105GF-Q3.3_07.15sec | 220 | FW | 100% |
| | | G07_RD14-0003-37_38_39_40-1_9_9_1-M2a-0.75GF-Q0.4_07.15sec | 0 | FW | 100% |
| | | H01_RD14-0003-40_41_42_43-1_1_1_1-M4d-0.124GF-Q1.7_08.15sec | 430 | SM OFF | 100% |
| | | H03_RD14-0003-44_45_46_47-1_1_4_1-M2e-0.441GF-Q1.5_08.15sec | 371 | BW & FW | 100% |
| θ = 0.03 | 2 | A02_RD14-0003-40_41-1_4-M3S30-0.075GF-Q4.0_01.15sec | 17 | SM OFF | 100% |
| | | A03_RD14-0003-40_41-1_4-M4a-0.075GF-Q0.6_01.15sec | 0 | SM OFF | 100% |
| | | B04_RD14-0003-40_41-1_4-M4e-0.075GF-Q0.7_02.15sec | 48 | BW & FW | 100% |
| | | D01_RD14-0003-44_45-1_1-M4I15-0.03GF-Q1.2_04.15sec | 254 | FW | 100% |
| | | E02_RD14-0003-31_32-1_1-M2d-0.03GF-Q6.0_05.15sec | 322 | SM OFF | 100% |
| | | F10_RD14-0003-39_40-1_2-M3c-0.045GF-Q1.0_06.15sec | 0 | SM OFF | 100% |
| | | G11_RD14-0003-39_40-1_2-M4e-0.045GF-Q2.4_07.15sec | 2 | SM OFF | 100% |
| | 3 | A02_RD14-0003-36_37_38-1_2_1-M4c-0.06GF-Q1.1_01.15sec | 44 | SM OFF | 100% |
| | | A08_RD14-0003-30_31_32-1_4_4-M2U60-0.135GF-Q2.7_01.15sec | 9 | SM OFF | 100% |
| | | A09_RD14-0003-49_50_29-1_4_1-M3e-0.186GF-Q1.6_01.15sec | 3 | SM OFF | 100% |
| | | B07_RD14-0003-49_50_29-1_4_1-M3U105-0.09GF-Q6.1_02.15sec | 315 | BW | 100% |
| | | B09_RD14-0003-49_50_29-1_4_1-M3e-0.09GF-Q1.6_02.15sec | 7 | SM OFF | 100% |
| | | D05_RD14-0003-46_47_48-1_1_1-M4I22-0.045GF-Q1.4_04.15sec | 353 | SM OFF | 100% |
| | | D08_RD14-0003-30_31_32-1_4_4-M2U105-0.135GF-Q5.1_04.15sec | 3 | SM OFF | 100% |
| | | F03_RD14-0003-49_50_29-1_4_1-M3S30-0.09GF-Q2.9_06.15sec | 352 | SM OFF | 100% |
| | | F04_RD14-0003-41_42_43-1_9_1-M2d-0.165GF-Q1.2_06.15sec | 334 | BW & FW | 100% |
| | | F09_RD14-0003-47_48_49-1_9_9-M2e-0.5GF-Q1.5_06.15sec | 21 | BW | 100% |
| | | H06_RD14-0003-49_50_29-1_4_1-M4I35-0.09GF-QLAND_08.15sec | 302 | FW | 100% |
| | 4 | A03_RD14-0003-50_29_30_31-1_1_2_1-M3e-0.075GF-Q1.1_01.15sec | 0 | SM OFF | 100% |
| | | C11_RD14-0003-48_49_50_29-1_4_4_4-M3d-0.195GF-Q0.9_03.15sec | 4 | SM OFF | 100% |
| | | D01_RD14-0003-40_41_42_43-1_1_1_1-M4I15-0.06GF-Q1.6_04.15sec | 42 | SM OFF | 100% |
| | | D09_RD14-0003-37_38_39_40-1_9_9_1-M2e-0.62GF-Q0.8_04.15sec | 25 | BW & FW | 100% |
| | | E01_RD14-0003-50_29_30_31-1_1_2_1-M3a-0.075GF-Q0.5_05.15sec | 0 | SM OFF | 100% |
| | | E04_RD14-0003-48_49_50_29-1_4_4_4-M2U105-0.195GF-Q12.1_05.15sec | 1 | BW | 100% |
| | | E05_RD14-0003-33_34_35_36-1_4_4_1-M2a-0.63GF-Q0.5_05.15sec | 0 | FW | 100% |
| | | E05_RD14-0003-36_37_38_39-1_2_2_1-M2e-0.378GF-Q1.2_05.15sec | 23 | BW | 100% |
| | | F05_RD14-0003-44_45_46_47-1_1_4_1-M2S30-0.217GF-Q3.3_06.15sec | 278 | SM OFF | 100% |
| | | G05_RD14-0003-44_45_46_47-1_1_4_1-M2S30-0.105GF-Q3.3_07.15sec | 193 | FW | 100% |
| | | G06_RD14-0003-33_34_35_36-1_4_4_1-M2d-0.15GF-Q0.9_07.15sec | 16 | FW | 100% |
| | | H02_RD14-0003-44_45_46_47-1_1_4_1-M3U60-0.105GF-Q28.7_08.15sec | 99 | BW | 100% |
