## Supplementary Figures 1-5 for "‘Low’ LRs obtained from DNA mixtures: On calibration and discrimination performance of probabilistic genotyping software"

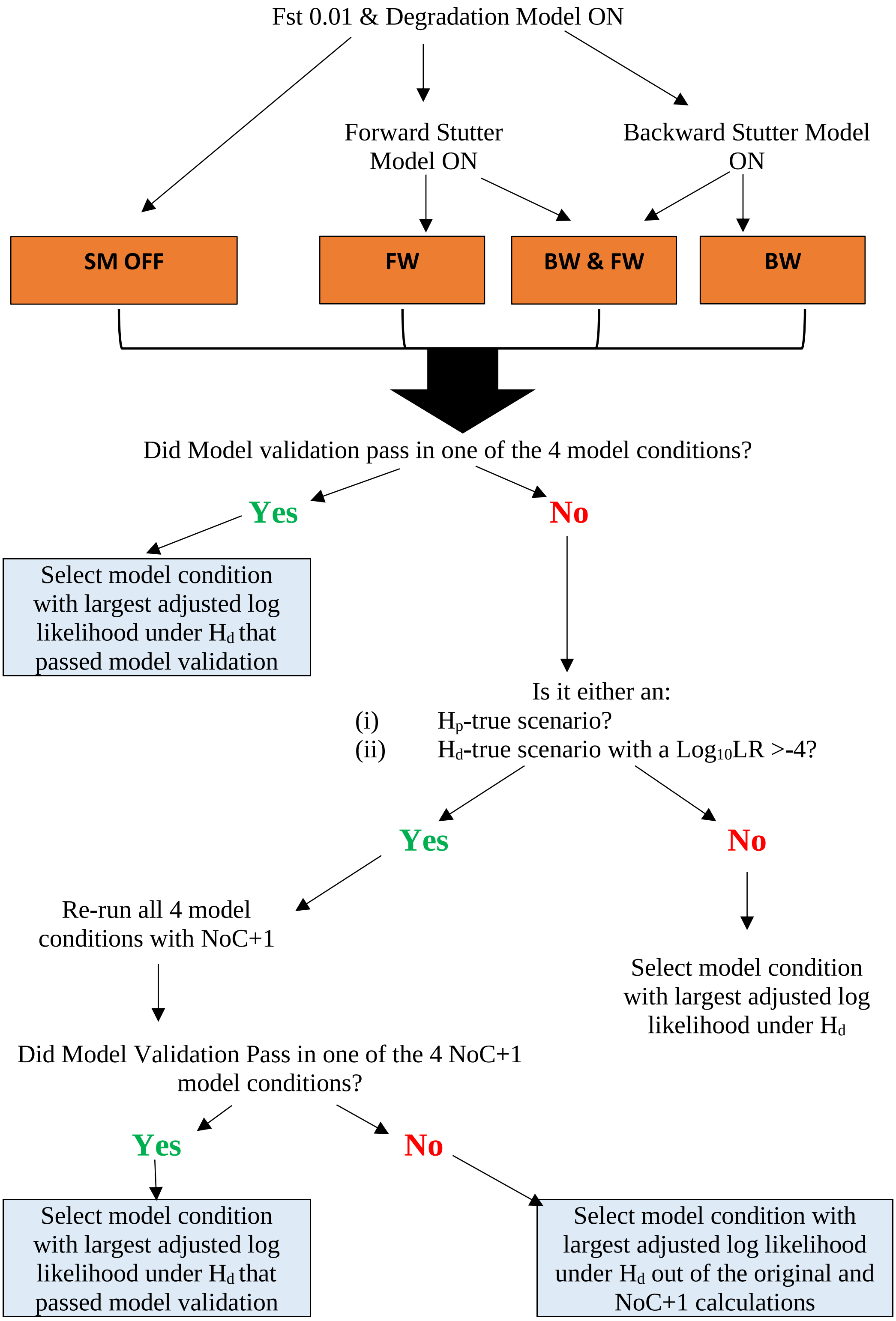


Supplementary Figure 1. DNAStatistX model selection process for each scenario. Note that blue boxes represent the final model selection step for a scenario.

HMC [17]

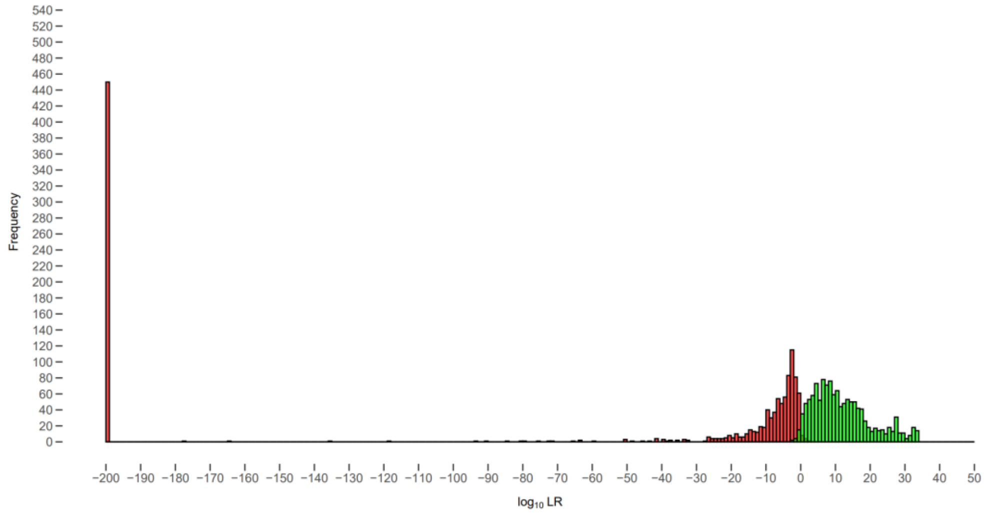


STRmix [17]

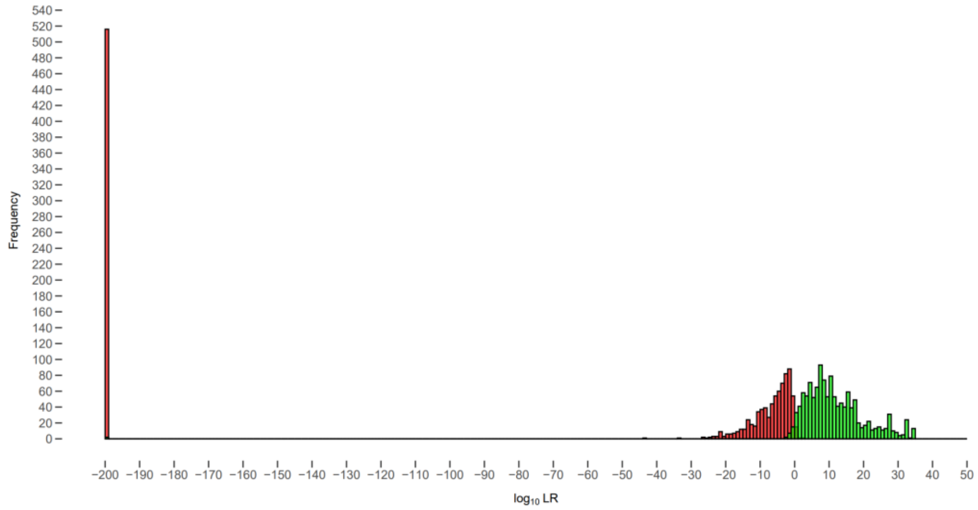


EFM

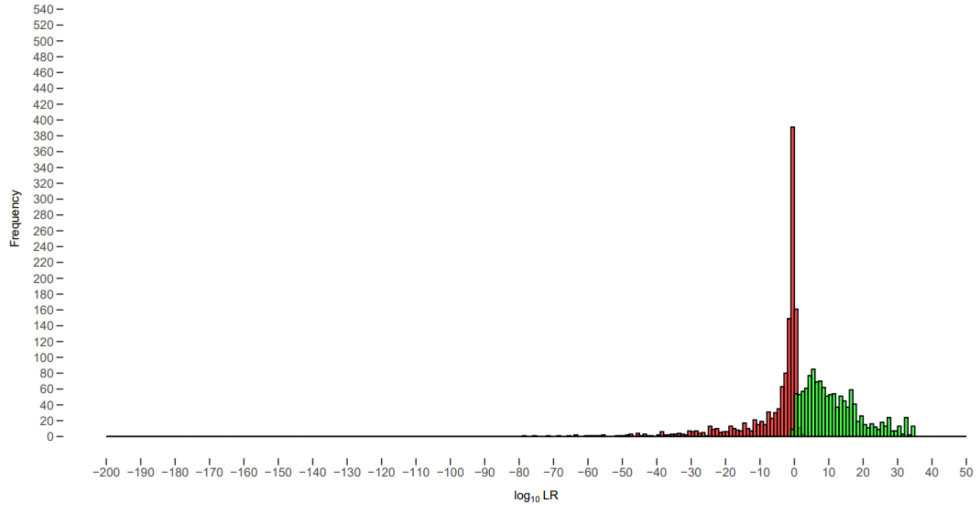

DNAStatistX

θ = 0.01

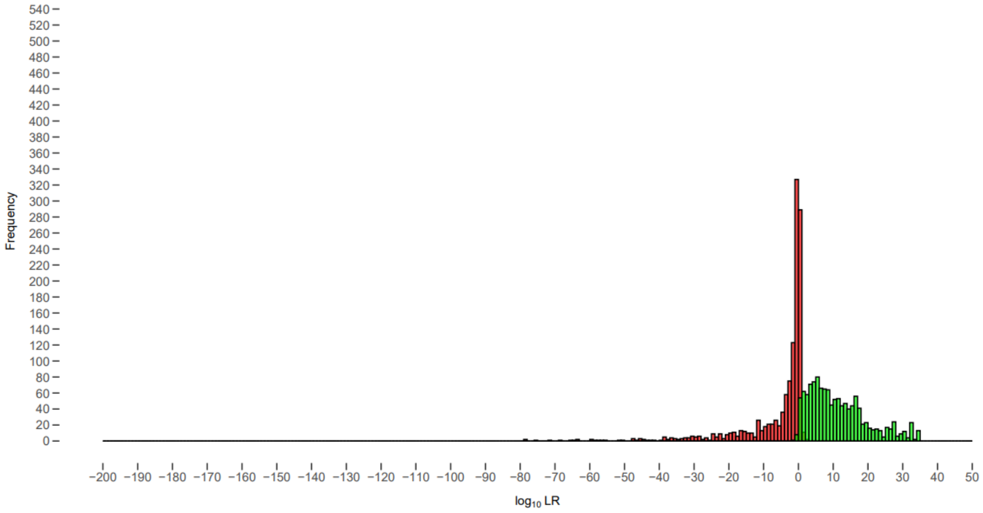


DNAStatistX

θ = 0.03

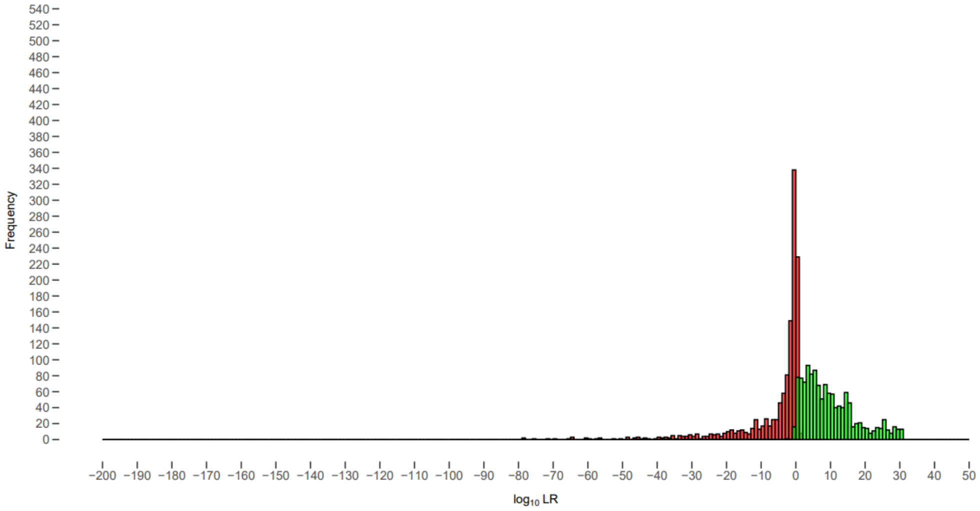


Supplementary Figure 2. Histograms of $\log_{10}$LRs of each PG software. Green: $H_p$-true scenarios, Red: $H_d$-true scenarios.

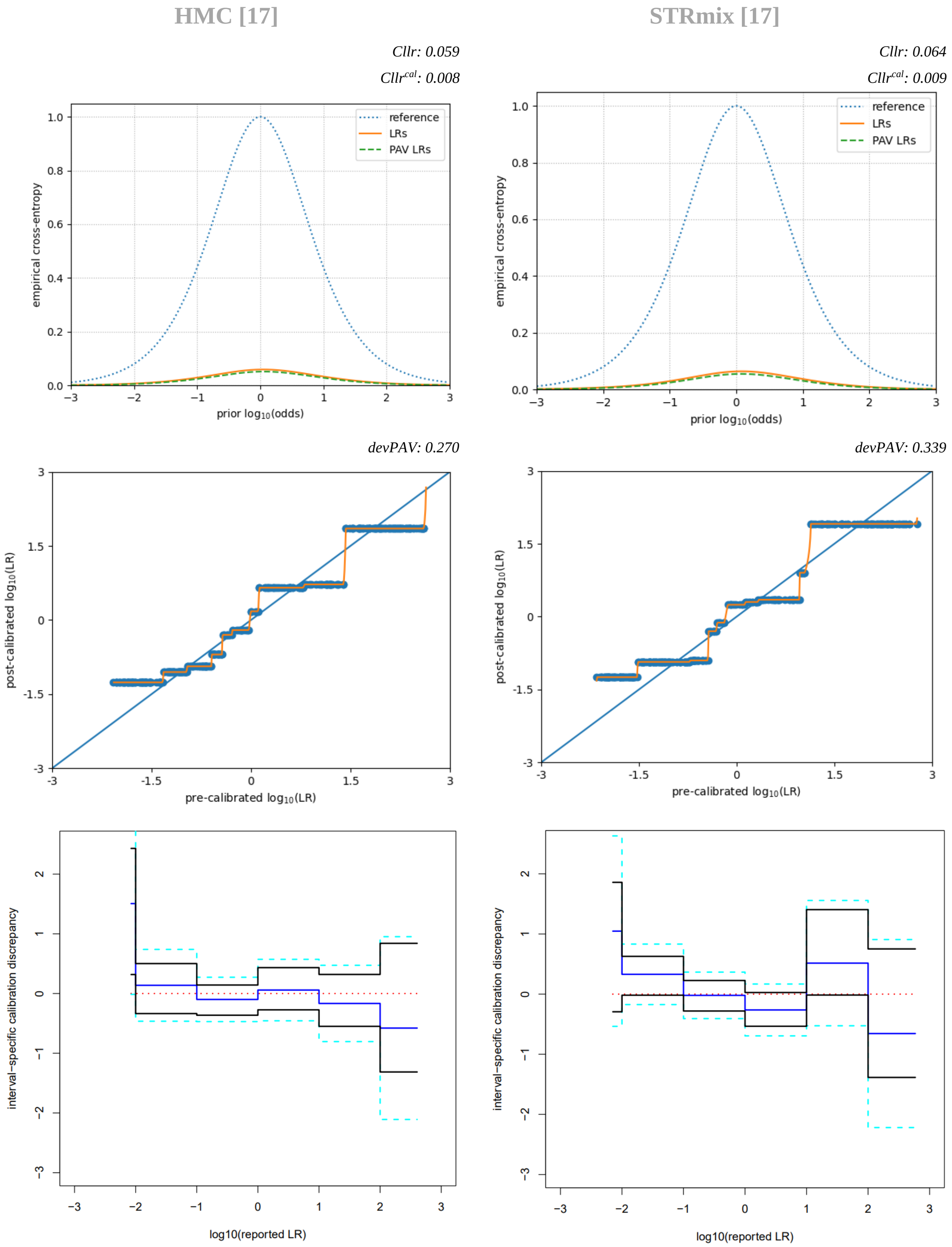


Supplementary Figure 3. Calibration plots of PG software. Top: ECE plots and Cllr values (3dp). Middle: PAV plots and devPAV values (3dp). Bottom: Fiducial calibration discrepancy plots.

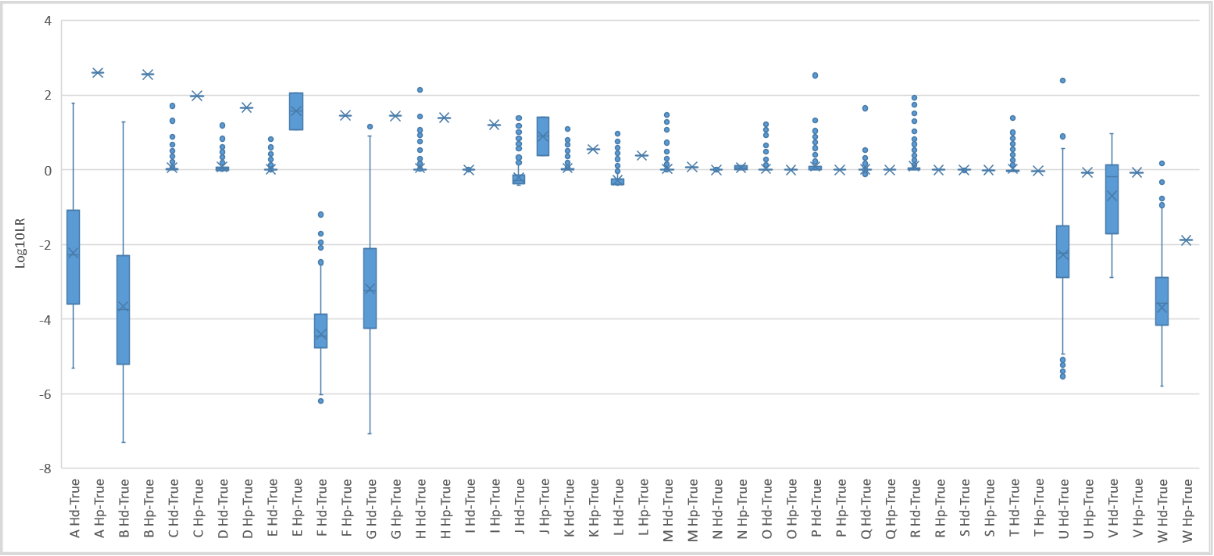


Supplementary Figure 4. Boxplots of $H_d$-true and $H_p$-true scenario $\log_{10}$LRs of DNAStatistX $\theta = 0.01$ in order of highest to lowest average $H_p$-true LR. Letters corresponding to scenarios summarised in the table below.

Letters corresponding to scenarios in Supplementary Figure 4.

| Mixture Profile | NoC | Selected Condition | Code |
|---|---|---|---|
| B04_RD14-0003-40_41-1_4-M4e-0.075GF-Q0.7_02.15sec | 2 | BW & FW | A |
| D03_RD14-0003-40_41-1_4-M3U105-0.075GF-Q14.8_04.15sec | 2 | SM OFF | B |
| H01_RD14-0003-40_41_42_43-1_1_1_1-M4d-0.124GF-Q1.7_08.15sec | 4 | SM OFF | C |
| E05_RD14-0003-36_37_38_39-1_2_2_1-M2e-0.378GF-Q1.2_05.15sec | 4 | BW | D |
| A03_RD14-0003-36_37_38-1_2_1-M3e-0.06GF-Q1.6_01.15sec | 3 | SM OFF | E |
| G07_RD14-0003-37_38_39_40-1_9_9_1-M2a-0.75GF-Q0.4_07.15sec | 4 | FW | F |
| G02_RD14-0003-31_32-1_1-M1e-0.062GF-Q14.4_07.15sec | 2 | SM OFF | G |
| H03_RD14-0003-44_45_46_47-1_1_4_1-M2e-0.441GF-Q1.5_08.15sec | 4 | BW & FW | H |
| D01_RD14-0003-44_45-1_1-M4I15-0.03GF-Q1.2_04.15sec | 2 | FW | I |
| C07_RD14-0003-33_34_35_36-1_4_4_1-M2e-0.15GF-Q0.9_03.15sec | 4 | SM OFF | J |
| G07_RD14-0003-30_31_32-1_4_4-M4I22-0.135GF-Q4.7_07.15sec | 3 | BW | K |
| F05_RD14-0003-46_47_48-1_1_1-M3U60-0.045GF-Q4.0_06.15sec | 3 | SM OFF | L |
| B05_RD14-0003-41_42_43-1_9_1-M2e-0.165GF-Q1.7_02.15sec | 3 | BW | M |
| H06_RD14-0003-49_50_29-1_4_1-M4I35-0.09GF-QLAND_08.15sec | 3 | FW | N |
| F05_RD14-0003-44_45_46_47-1_1_4_1-M2S30-0.217GF-Q3.3_06.15sec | 4 | SM OFF | O |
| G05_RD14-0003-44_45_46_47-1_1_4_1-M2S30-0.105GF-Q3.3_07.15sec | 4 | FW | P |
| B07_RD14-0003-49_50_29-1_4_1-M3U105-0.09GF-Q6.1_02.15sec | 3 | BW | Q |
| F03_RD14-0003-49_50_29-1_4_1-M3S30-0.09GF-Q2.9_06.15sec | 3 | SM OFF | R |
| G11_RD14-0003-39_40-1_2-M4e-0.045GF-Q2.4_07.15sec | 2 | SM OFF | S |
| A09_RD14-0003-49_50_29-1_4_1-M3e-0.186GF-Q1.6_01.15sec | 3 | SM OFF | T |
| D08_RD14-0003-30_31_32-1_4_4-M2U105-0.135GF-Q5.1_04.15sec | 3 | SM OFF | U |
| F10_RD14-0003-39_40-1_2-M3c-0.045GF-Q1.0_06.15sec | 2 | SM OFF | V |
| E05_RD14-0003-33_34_35_36-1_4_4_1-M2a-0.63GF-Q0.5_05.15sec | 4 | FW | W |

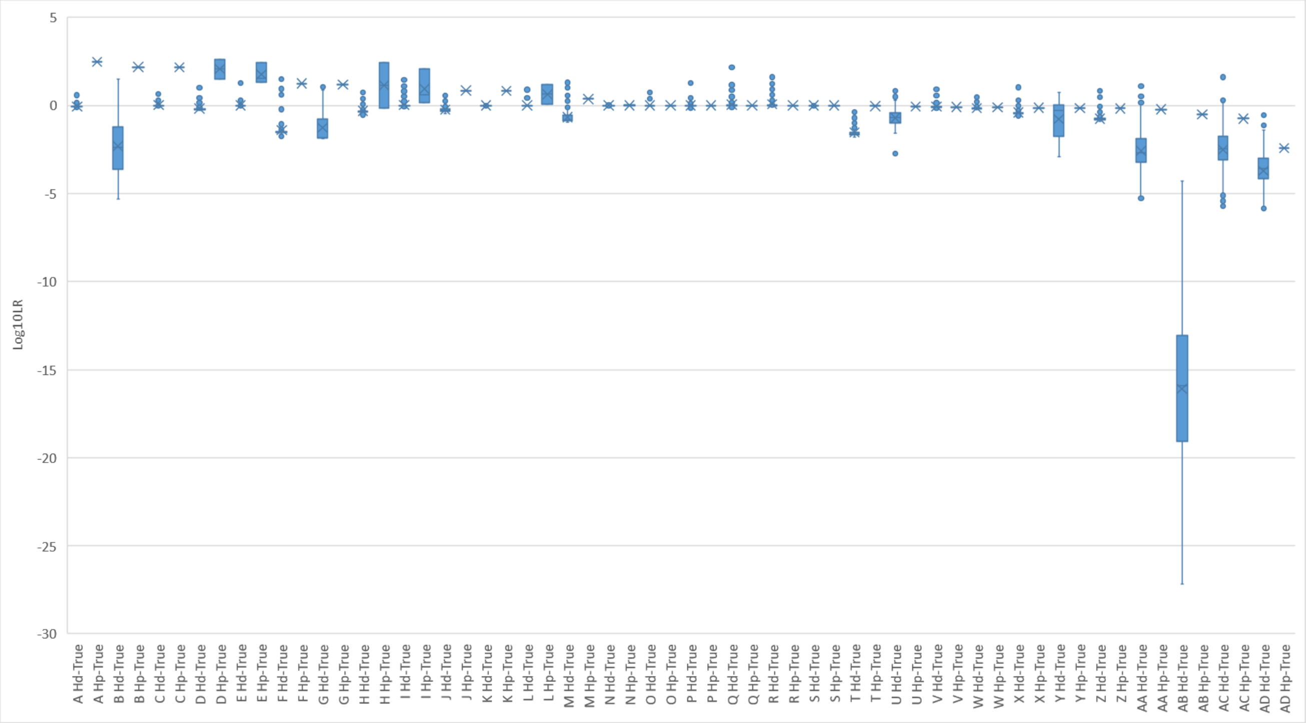


Supplementary Figure 5. Boxplots of $H_d$-true and $H_p$-true scenario $log_{10}$LRs of DNAStatistX θ = 0.03 in order of highest to lowest average $H_p$-true LR. Letters corresponding to scenarios summarised in the table below.

Letters corresponding to scenarios in Supplementary Figure 5.

| Mixture Profile | NoC | Selected Condition | Code |
|---|---|---|---|
| D01_RD14-0003-40_41_42_43-1_1_1_1-M4I15-0.06GF-Q1.6_04.15sec | 4 | SM OFF | A |
| B04_RD14-0003-40_41-1_4-M4e-0.075GF-Q0.7_02.15sec | 2 | BW & FW | B |
| F04_RD14-0003-41_42_43-1_9_1-M2d-0.165GF-Q1.2_06.15sec | 3 | BW & FW | C |
| A02_RD14-0003-36_37_38-1_2_1-M4c-0.06GF-Q1.1_01.15sec | 3 | SM OFF | D |
| D05_RD14-0003-46_47_48-1_1_1-M4I22-0.045GF-Q1.4_04.15sec | 3 | SM OFF | E |
| F09_RD14-0003-47_48_49-1_9_9-M2e-0.5GF-Q1.5_06.15sec | 3 | BW | F |
| A02_RD14-0003-40_41-1_4-M3S30-0.075GF-Q4.0_01.15sec | 2 | SM OFF | G |
| G06_RD14-0003-33_34_35_36-1_4_4_1-M2d-0.15GF-Q0.9_07.15sec | 4 | FW | H |
| H02_RD14-0003-44_45_46_47-1_1_4_1-M3U60-0.105GF-Q28.7_08.15sec | 4 | BW | I |
| E05_RD14-0003-36_37_38_39-1_2_2_1-M2e-0.378GF-Q1.2_05.15sec | 4 | BW | J |
| D01_RD14-0003-44_45-1_1-M4I15-0.03GF-Q1.2_04.15sec | 2 | FW | K |
| E02_RD14-0003-31_32-1_1-M2d-0.03GF-Q6.0_05.15sec | 2 | SM OFF | L |
| D09_RD14-0003-37_38_39_40-1_9_9_1-M2e-0.62GF-Q0.8_04.15sec | 4 | BW & FW | M |
| H06_RD14-0003-49_50_29-1_4_1-M4I35-0.09GF-QLAND_08.15sec | 3 | FW | N |
| F05_RD14-0003-44_45_46_47-1_1_4_1-M2S30-0.217GF-Q3.3_06.15sec | 4 | SM OFF | O |
| B07_RD14-0003-49_50_29-1_4_1-M3U105-0.09GF-Q6.1_02.15sec | 3 | BW | P |
| G05_RD14-0003-44_45_46_47-1_1_4_1-M2S30-0.105GF-Q3.3_07.15sec | 4 | FW | Q |
| F03_RD14-0003-49_50_29-1_4_1-M3S30-0.09GF-Q2.9_06.15sec | 3 | SM OFF | R |
| G11_RD14-0003-39_40-1_2-M4e-0.045GF-Q2.4_07.15sec | 2 | SM OFF | S |
| E01_RD14-0003-50_29_30_31-1_1_2_1-M3a-0.075GF-Q0.5_05.15sec | 4 | SM OFF | T |
| B09_RD14-0003-49_50_29-1_4_1-M3e-0.09GF-Q1.6_02.15sec | 3 | SM OFF | U |
| A09_RD14-0003-49_50_29-1_4_1-M3e-0.186GF-Q1.6_01.15sec | 3 | SM OFF | V |
| A03_RD14-0003-50_29_30_31-1_1_2_1-M3e-0.075GF-Q1.1_01.15sec | 4 | SM OFF | W |
| E04_RD14-0003-48_49_50_29-1_4_4_4-M2U105-0.195GF-Q12.1_05.15sec | 4 | BW | X |
| F10_RD14-0003-39_40-1_2-M3c-0.045GF-Q1.0_06.15sec | 2 | SM OFF | Y |
| C11_RD14-0003-48_49_50_29-1_4_4_4-M3d-0.195GF-Q0.9_03.15sec | 4 | SM OFF | Z |
| A08_RD14-0003-30_31_32-1_4_4-M2U60-0.135GF-Q2.7_01.15sec | 3 | SM OFF | AA |
| A03_RD14-0003-40_41-1_4-M4a-0.075GF-Q0.6_01.15sec | 2 | SM OFF | AB |
| D08_RD14-0003-30_31_32-1_4_4-M2U105-0.135GF-Q5.1_04.15sec | 3 | SM OFF | AC |
| E05_RD14-0003-33_34_35_36-1_4_4_1-M2a-0.63GF-Q0.5_05.15sec | 4 | FW | AD |
